# Discovery of a hidden transient state in all bromodomain families

**DOI:** 10.1101/2020.04.01.019547

**Authors:** Lluís Raich, Katharina Meier, Judith Günther, Clara D. Christ, Frank Noé, Simon Olsson

**Author notes:** Correspondence: Simon Olsson or Frank Noé.

## Abstract

Bromodomains (BDs) are small protein modules that interact with acetylated marks in histones. These post-translational modifications are pivotal to regulate gene expression, making BDs promising targets to treat several diseases. While the general structure of BDs is well known, their dynamical features and their interplay with other macromolecules are poorly understood, hampering the rational design of potent and selective inhibitors. Here we combine extensive molecular dynamics simulations, Markov state modeling and structural data to reveal a novel and transiently formed state that is conserved across all BD families. It involves the breaking of two backbone hydrogen bonds that anchor the ZA-loop with the α_A_ helix, opening a cryptic pocket that partially occludes the one associated with histone binding. Our results suggest that this novel state is an allosteric regulatory switch for BDs, potentially related to a recently unveiled BD-DNA binding mode.

## INTRODUCTION

Proteins are highly dynamic biomolecules, often adopting multiple conformational states that can be critical to their function, regulation and evolvability.^1–3^ Transitions between the different conformations of a protein may directly connect to changes in several properties, such as protein stability, interactions with binding-partners, or the appearance of “cryptic” pockets that could be druggable but are hidden in native structures.^4–6^ Characterizing and determining such conformations is therefore essential to understanding the intricate relationship between structure, dynamics and function of proteins, with a direct impact on drug design and protein engineering.^7,8^ Unfortunately, most of the conformations that a protein can adopt have low populations, and are consequently difficult to characterize with conventional biophysical techniques. Indeed, minor conformational states may be masked under certain experimental conditions, making their detection challenging.^9,10^ Nevertheless, several methods have emerged to overcome these limitations, including ambient-temperature X-ray crystallography,^11,12^ NMR relaxation dispersion experiments,^13,14^ and extensive computer simulations,^15,16^ all of which have been successful in the discovery of such minor-but-relevant states.

Human bromodomains (BDs) are important epigenetic regulators, recognizing or “reading” acetylation marks in histone tails.^17^ These domains are part of larger proteins that may also include enzymatic domains to modify chromatin structure, or binding domains that can serve as platforms to recruit and activate transcription factors.^18^ Their central role make BD-containing proteins pivotal to regulate gene expression, representing crucial targets for the treatment of several diseases.^19^ Significant efforts have gone into characterizing the structure of BDs, aiming to discern the differences between the eight BD families. These studies revealed that all BDs share a common fold comprised by a 4-helix bundle (α_Z_,α_A_,α_B_,α_C_) with two principal loops that connect them (ZA and BC loops), conforming a hydrophobic pocket that is suited for acetyl-lysine binding (see Figure 1).^20^ The BC loop contains a well-conserved hydrogen bond donor or “reader” –usually an asparagine– that is key for recognition. The ZA-loop is the longest loop and the one that displays more variability in terms of sequence and structure, presenting short helices, turns and even hairpin insertions in BD Family VIII (Fig. 1). This loop is anchored to the α_A_ helix via two conserved backbone hydrogen bonds (h-bonds), forming a cavity that is known as the ZA channel, an important structural feature for optimizing the potency and selectivity of BD inhibitors.^19,21^ Characterization of BD flexibility has been, on the other hand, limited to a few studies reporting different crystallographic structures and molecular dynamics (MD) simulations that show subtle motions of the ZA-loop, as well as transitions between rotameric states of residues involved in ligand binding.^22–24^

**Figure 1.**
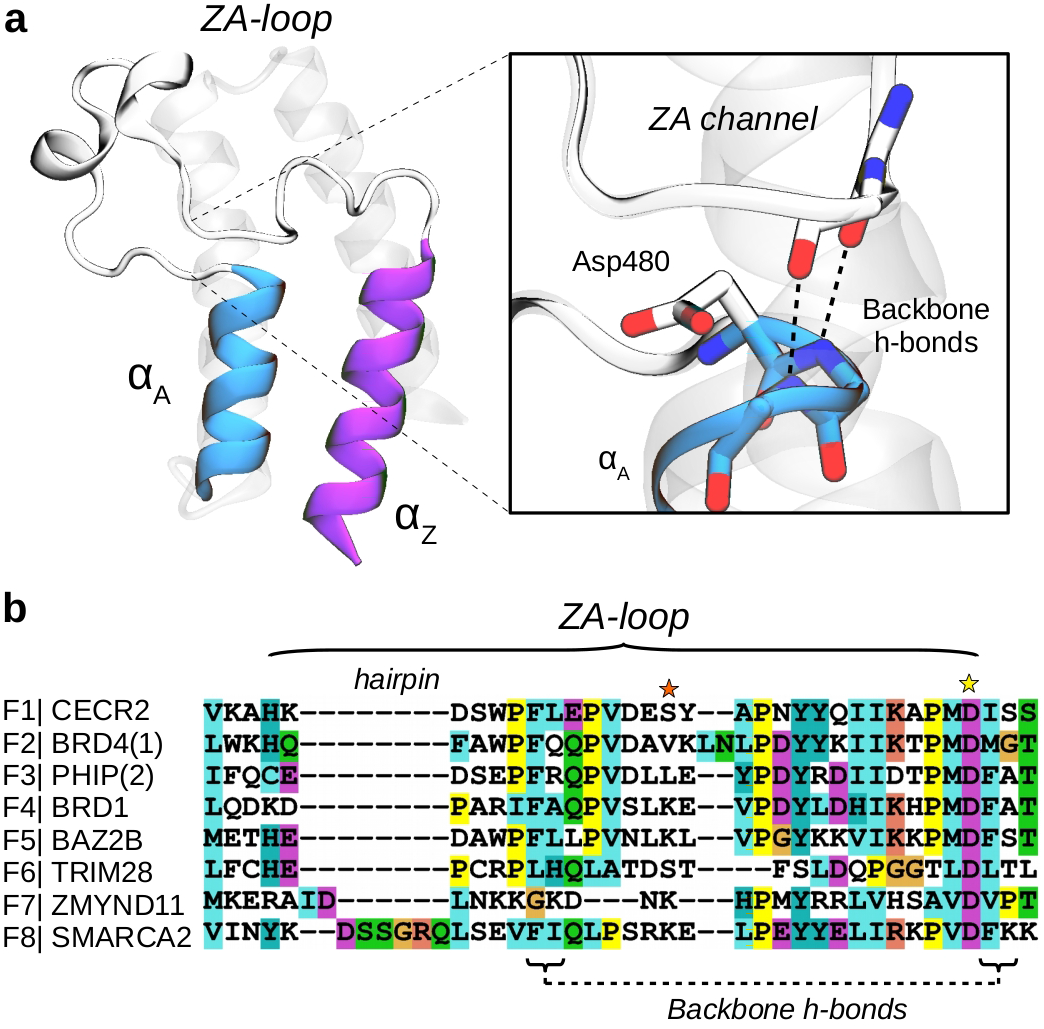
Bromodomains have two conserved backbone h-bonds that anchor the ZA-loop with the α_A_ helix. (**a**) Structure of CECR2 (PDB 3NXB) highlighting the two first helices, αZ and αA, and the long ZA-loop, detailing the ZA channel and the two conserved backbone h-bonds that anchor the loop with the αA helix. (**b**) Sequence alignment of the ZA-loop region for the BDs analyzed in this work, including all families. Note the presence of a hairpin insertion in family VIII, and the well conserved Asp480 (yellow star). The orange star highlights lysine residues commented below.

Although these insights have helped in the general understanding of BDs, the poor comprehension of their conformational ensembles, their biological relevance within BD-containing proteins and their precise role in disease are hampering the advance of BD inhibitors into the clinic.^25^ A recent study illustrates this challenge through a detailed characterization of the impact of missense variation on BD function.^26^ The authors show that while specific variants did not alter the global fold of BDs in crystal structures, spectroscopic measurements in solution suggest changes in dynamic properties that could account for the loss of protein stability and inhibitor affinity. Furthermore, another recent study demonstrated that BDs can directly bind to DNA through a basic patch on their surface and that in fact DNA-binding drives the association of BDs with chromatin.^27^ These macroscopic interactions could affect the structure and flexibility of BDs, modulating their function and properties, which opens questions on the relevance and precise role of BDs inside their biological context.

MD simulations have been pivotal to complement experiments in the understanding of protein function, focusing on their dynamics and flexibility. MD simulations, in principle, grant direct access to proteins at the full spatial and temporal resolution. However, extensive simulations may be needed to sample the rare transitions between long-lived protein conformations. The combination of high-throughput MD simulations with Markov state models (MSMs) provides a powerful framework to recover long-time dynamics from multiple short-time simulations, following a “divide-and-conquer” philosophy.^28^ These techniques have enabled the quantitative study of critical biological processes such as protein folding or protein-protein association,^29,30^ protein plasticity and ligand-binding kinetics,^31^ as well as to characterize cryptic pockets in proteins and find common states across protein families,^8,32^ making them very useful to evaluate the structural flexibility of BDs.

Here, using MD simulations and MSMs, we survey representative BDs from the eight families to characterize their conformational dynamics. Although the analyzed proteins share little sequence identity, we identify two recurring, transient and novel states in which the two backbone hydrogen bonds that anchor the ZA-loop with the α_A_ helix break, and the acetyl-lysine binding pocket becomes occluded. The population of this new state is generally smaller than that of the crystallographic state, suggesting why so far it has escaped experimental observation. We propose that this conformational change represents a potential mechanism of auto-regulation in BDs that is related to a BD-DNA binding mode, and reveal allosteric pockets that could be leveraged to modulate both histone and DNA binding contributions to chromatin, offering new starting points to overcome the current challenges facing the design of potent and selective BD inhibitors.

## RESULTS

### All BD families share a conserved and novel metastable state

We started our study focusing on BRD4(1), a thoroughly characterized bromodomain (BD) of family II. We ran 64 independent 1 microsecond MD simulations to build a MSM using the PyEMMA software (see Supplementary methods for details).^33^ Analysis of the resulting model reveals a novel metastable state apart from the one that resembles the crystallographic structure, which involves the displacement of the ZA-loop from the α_A_ helix, opening a space beneath it that increases the solvent accessible surface area of the well-conserved aspartate (Asp106, see Figure 2). Interestingly, the conformational change disrupts the ZA channel, a structural feature that is relevant for inhibitor selectivity. In terms of interactions, the opening process involves the breaking of the two conserved backbone h-bonds, whose interactions are partially compensated by the h-bonds that Gln84 establishes with Asp106, acting as a “latch”. The free energy profile along the slowest time-lagged independent component (TIC^34,35^) shows two clear basins, with the open state being ~2 kcal·mol^−1^ above the closed (Figure 2c). The mean first passage time (MFPT) for the opening is of 5.9 ± 1.2 μs and 688 ± 61 ns for the closing, indicated by medians and one standard deviation from a bootstrapping distribution.

**Figure 2.**
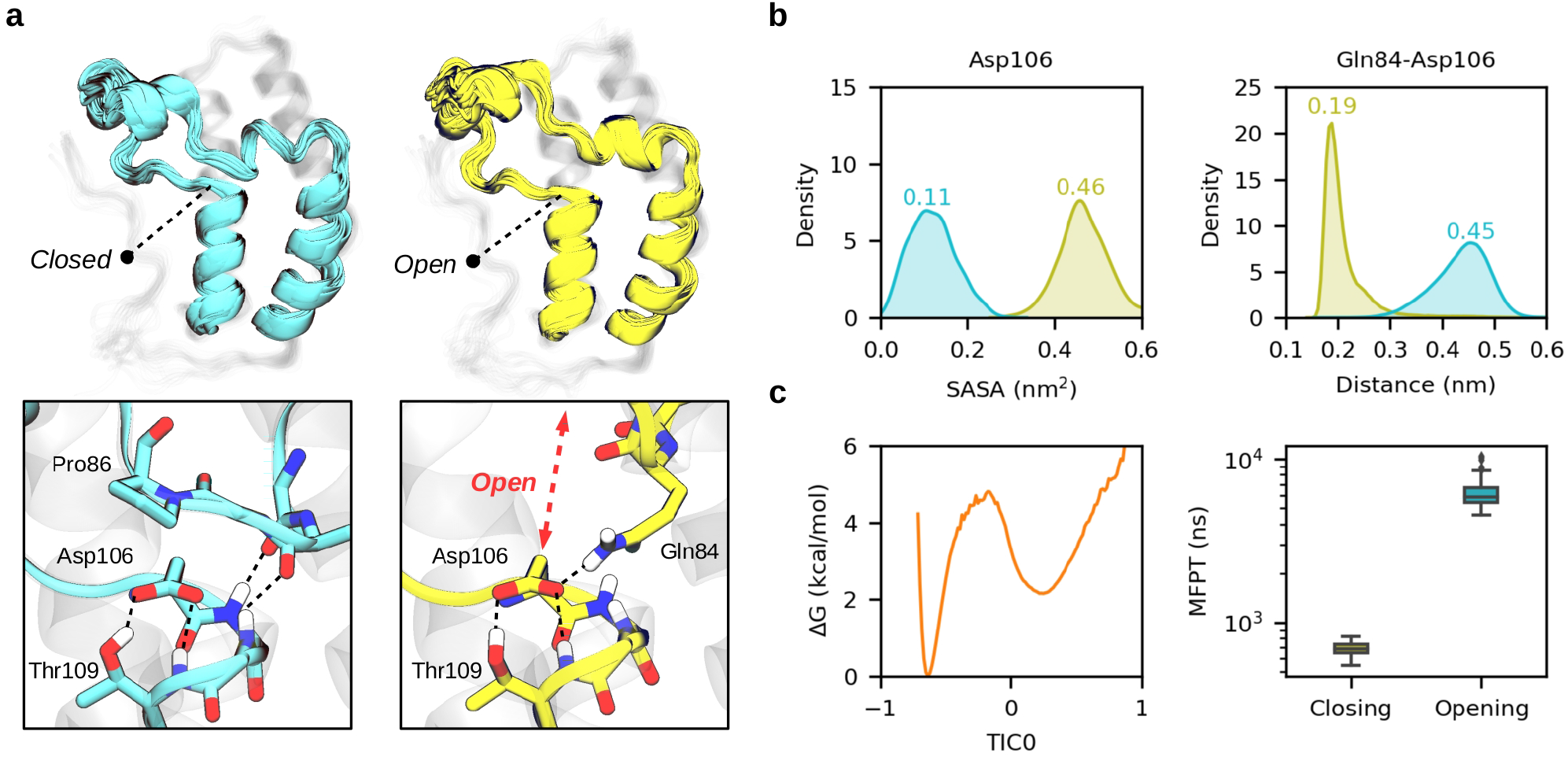
A novel conformational state in BDs. (**a**) Ensemble of structures of the “closed” crystal-like state (cyan) and the “open” novel state (yellow) for BRD4(1). A close view of the opening region is represented below, highlighting Asp106. (**b**) Distribution of the Asp106 solvent accessible surface area (SASA) and closest distance between Gln84-Asp106 side chains for the two metastable states. (**c**) MSM reweighted free energy profile along the first independent component (TIC0, arbitrary units) and boxplot of a bootstrapped distribution of the closing and opening mean first passage times.

The fact that the opening implies the breaking of two backbone h-bonds, likely present in all BDs, prompted us to analyze the structural flexibility of other BD families, wondering whether this could be a general conformational feature across them. We selected a member of each BD family based on a sequence alignment, focusing on a set of residues that lay around the conserved aspartate that is beneath the ZA-loop. In total, ordered by families, we decided to study CECR2, BRD4(1), PHIP(2), BRD1, BAZ2B, TRIM28, ZMYND11 and SMARCA2 (see Supplementary Table 1 for the PDB identifiers and amino acid sequences). We ran 40 to 60 independent 1 μs MD for each BD, collecting an aggregate simulation data of 392 μs, which represents, as far as we know, the most extensive computational study of BDs.

Strikingly, our results reveal that the novel state is conserved in all families, with slight differences in their populations, exchange timescales, and structural features (see Figure 3, Supplementary Figures 2-9, and Supplementary Tables 2-3). For instance, CECR2 and PHIP(2) show a population of the open state above 30%, BRD4(1), BRD1 and BAZ2B around 10%, and SMARCA2 below 5%. ZMYND11 is a special case because it is always in the open state –*i.e*. 100%– even though we observe a “semiclosed” state in which the extreme part of the ZA-loop partially closes. TRIM28 is another special case, as its ZA-loop is wrapped on top of the α_A_ and α_B_ helices in the experimental structure. It presents a quite unstable fold and a complex conformational landscape that would require an extensive simulation effort to compute state populations and rates reliably, and thus we do not provide values in this case. Nonetheless, we have observed several trajectories showing clear metastability in the open state, confirming its kinetic relevance (Supplementary Figure 7). In terms of timescales, CECR2 is the BD that opens fastest, with a median MFPT of 182 ± 7 ns, followed by PHIP(2) with 903 ± 70 ns, BAZ2B 2.6 ± 0.5 μs, BRD4(1) 5.9 ± 1.2 μs, BRD1 8.3 ± 4.1 μs and SMARCA2 11.5 ± 1.5 μs (see Supplementary Table 3). The closing event is generally one order of magnitude faster than the opening, all of them below 1 μs, having SMARCA2 closing within 68 ± 3 ns, followed by CECR2 110 ± 4 ns, BAZ2B 377 ± 57 ns, PHIP(2) 449 ± 51 ns, BRD1 643 ± 117 ns and BRD4(1) 688 ± 61 ns. These results may explain why the novel state has been elusive in previous studies, since published simulation timescales are usually around few hundreds of nanoseconds,^36–40^ which is insufficient to characterize the opening event for most of the BDs analyzed here.

**Figure 3.**
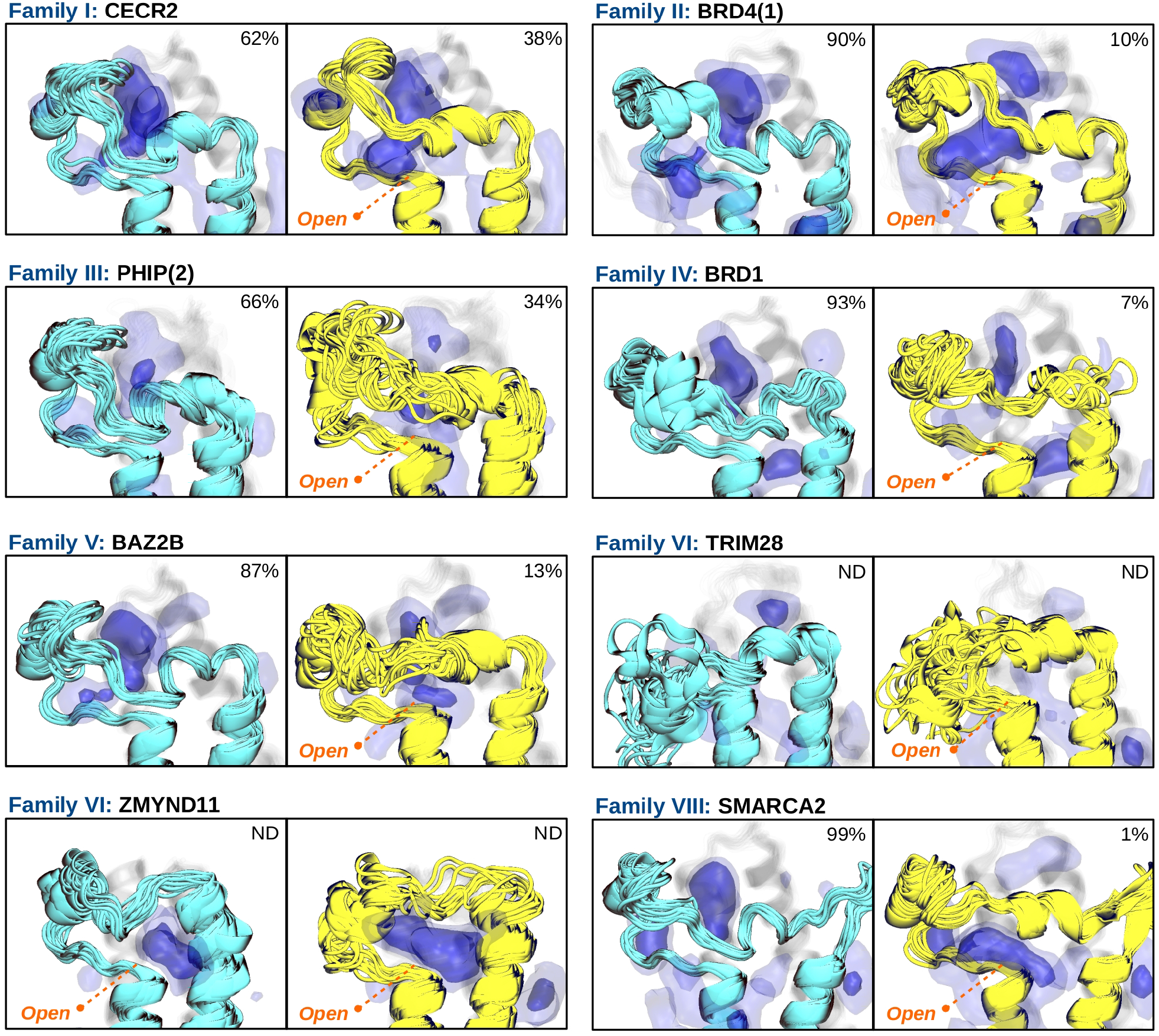
All bromodomain families share a novel state that opens a cryptic pocket beneath the ZA-loop. Ensemble of structures of the “closed” crystal-like state (cyan) and the “open” novel state (yellow) for each BD family. MDpocket frequency maps are represented by blue isosurfaces at 0.25 (light) and 0.50 (intense) isovalues, highlighting different binding sites. Percentages refer to medians of MSM populations obtained from a bootstrapped distribution (Supplementary Table 2). Populations for TRIM28 have not been determined due to its complex conformational landscape, and neither the ones of ZMYND11 since all structures are in the open state.

At a structural level, the novel state shows an overall similarity between BDs, with a clear displacement of the ZA-loop from the α_A_ helix (Figure 3). Although this conformational change generally involves the breaking of the two backbone h-bonds (see Supplementary Table 4 and Supplementary Figures 10-11), there are subtle differences between systems that are worth noting. For instance, CECR2 is the only BD that retains a substantial percentage of h-bond 1 in the open state, and thus the opening process is better described by the breaking of h-bond 2. BRD1 is another interesting system in this regard, since the two key h-bonds are only transiently formed in the closed state. This happens as BRD1 switches between the crystallographic state and an alternative state in which the ZA-loop is slightly displaced towards the α_Z_ helix, breaking the h-bonds (see Supplementary Fig. 12). The loss of these key interactions is compensated by backbone amides of the ZA-loop, establishing h-bonds with the conserved aspartate to stabilize the closed state, resembling the interactions occurring in TRIM28 and in most BDs of Family VIII (see Supplementary Figs. 7a and 9a). Interestingly, this alternative state is also present in CECR2, PHIP(2) and BAZ2B, albeit much less populated. Finally, we note that the majority of the BDs analyzed here display sporadic opening events in which either one or the two h-bonds break. These openings occur within a microsecond timescale and are generally not metastable (Supplementary Figs. 13-14). This observation suggests a duality between the classical mechanisms of “fast” induce-fit and “slow” conformational selection, both of which may be interesting for drug design.

### A cryptic and allosteric pocket opens in the novel state

An intriguing aspect of the conformational change is that it always disrupts the ZA channel, reducing the accessibility to the acetyl-lysine pocket (Supplementary Fig. 15). This is relevant since the ZA channel is a key structural element for the design of inhibitors, conferring them potency and selectivity.^41–43^ Therefore, finding pockets whose targeting could stabilize the open state would be suitable for controlling BD function. To that end, we have used the MDpocket software to analyze a thousand representative structures of each metastable state for every BD.^44^ The results unequivocally show the presence of novel binding pockets in the open state, just beneath the ZA-loop (Fig. 3). We classify these cryptic pockets in two groups depending on their opening region. For instance, in CECR2, BRD4(1), BRD1 and SMARCA2 the pockets mostly open in the region where the ZA-loop was lining, next to the well-conserved aspartate, while in BAZ2B and ZMYND11 the pockets open more in between the ZA-loop and the α_Z_ and α_A_ helices. We attribute this difference to a lysine residue found in the upper part of the ZA-loop (Fig. 1b, orange star), that reorients to interact with the exposed aspartate (see Supplementary Figures 6 and 8). We note that BRD1 also has an alternative conformational state, not included in the model, in which the pocket is displaced towards the α_Z_ and α_A_ helices by a lysine residue (Supplementary Fig. 16). Interestingly, the presence of this lysine in SMARCA2 does not have the same effect, which may be due to the electrostatic shielding of two charged residues that are just at the beginning of the α_A_ helix, flanking the conserved aspartate (Supplementary Fig. 9).

The binding sites of PHIP(2) are less defined, both in the open and closed states, given the large fluctuations of the ZA-loop, which partially obstructs the acetyl-lysine pocket. However, relying on the opening motion and the lack of a pocket in the central part, we can include this BD in the first group together with CECR2, BRD4(1), BRD1 and SMARCA2. Interestingly, the partial obstruction of the acetyl-lysine pocket is in agreement with the fact that PHIP(2) has been difficult to target, and thus only a recent study reports a few hits that weakly bind the outermost region of the pocket.^45^ These results highlight the importance of considering structural ensembles instead of single structures to determine the druggability of a pocket.^46^

TRIM28 is an exception in terms of pockets given the unusual conformation of the ZA-loop. In the closed state, for instance, it increases the solvent exposure of the acetyl-lysine pocket such that it is not anymore defined, and in the open state it fluctuates so much that no pockets appear in the region of interest.

Finally, we have superposed the closed state of our models with crystal structures that have drugs bound in the acetyl-lysine pocket, finding an excellent agreement between the pose of all drugs and the computed isosurfaces (see Supplementary Figures 2-6 and 9), including a deeply buried drug in SMARCA2, for which a set of conserved water molecules of the pocket are known to be displaceable.^43,47^

### Crystallographic evidence confirms the existence of the open state

To validate our results, we analyzed all the structures that are present in the bromodomain entry (PF00439) of the Pfam database,^48^ accounting a total of 1.903 structures (see *online materials* for the complete list of PDBs). We have computed the distances of the two backbone h-bonds for all structures, and projected them onto the free energy landscape of BRD4(1) obtained with the same descriptors. Remarkably, the experimental distribution of h-bonds closely overlaps with the low free energy regions of the landscape, showing a small variance for the first h-bond and a more pronounced variance for the second (see Fig. 4a and Supplementary Fig. 17 for the projection onto the other BDs). This is a surprising result since the analyzed structures are of BDs with very different sequences, including organisms that are genetically far from humans, and yet the free energy landscape of a single BD encloses all this variation. Most importantly, this analysis has allowed us to detect outliers in the distribution, and find two crystal structures that are stable in the open state (Fig. 4b,c). These structures are the ones of ZMYND11 and PB1(6), BDs that present substantial sequence variation –particularly in the ZA-loop region– compared to the general trend.

**Figure 4.**
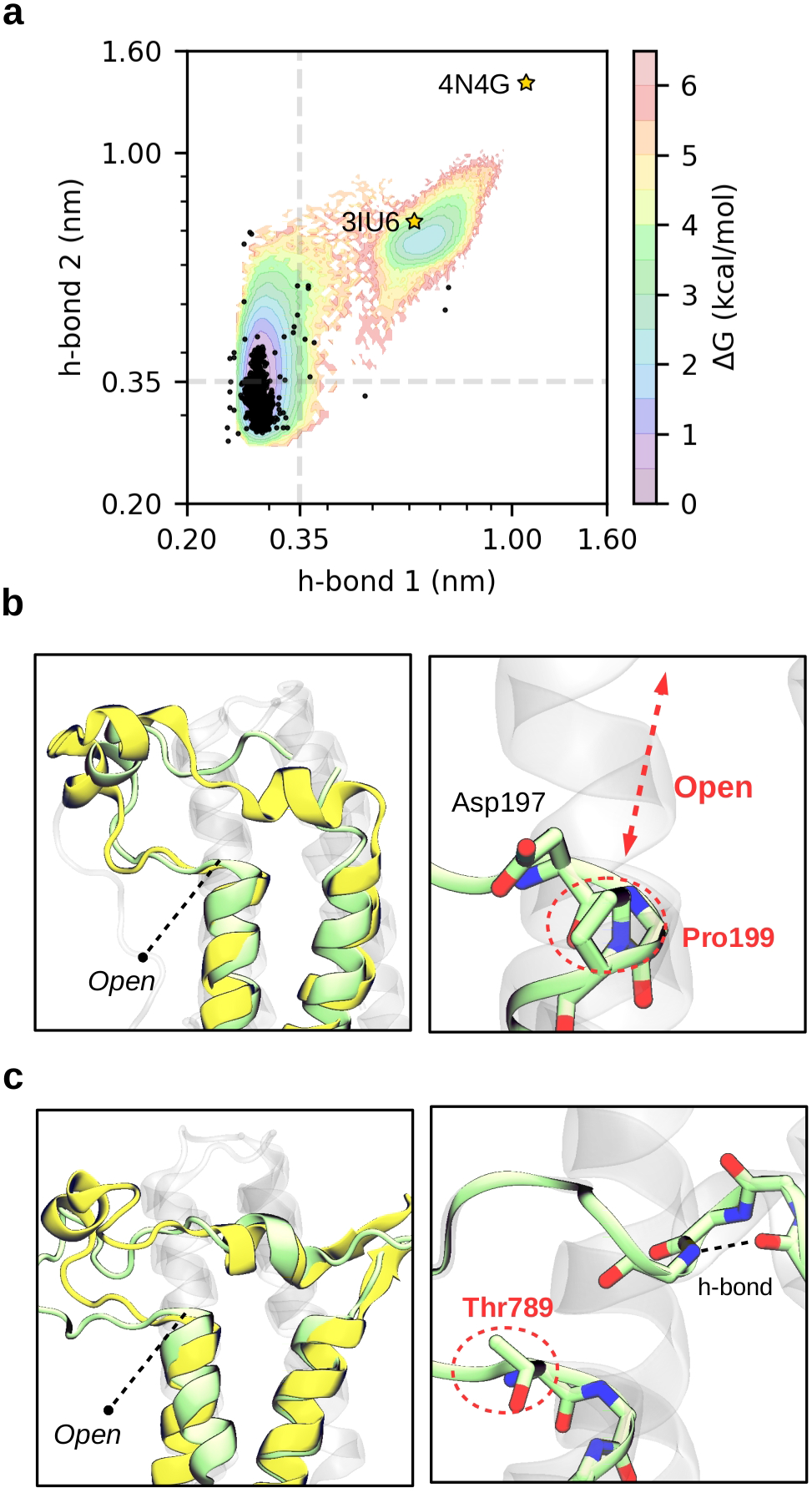
Distribution of h-bonds in experimentally determined structures reveal two BDs in the open state. (**a**) Projection of all BD structures from the Pfam database (black dots) on a MSM reweighted free energy landscape of BRD4(1) comprising the two conserved backbone h-bonds. Axes are given in a logarithmic scale and dashed lines indicate a distance of 0.35 nm as an upper bound for h-bond formation. The stars highlight the two crystal structures that are in the open state. (**b**) The structure of ZMYND11 (pale green, PDB 4N4G) is compared with the open state predicted for BRD4(1) (yellow). Pro199 is highlighted next to the conserved Asp. (**c**) The structure of PB1(6) (pale green, PDB 3IU6) is compared with the open state predicted for SMARCA2 (yellow). Thr789, in place of the conserved Asp, is highlighted together with an internal h-bond that is formed in the short helix of the ZA-loop.

A close inspection of ZMYND11 reveals the presence of a proline residue (Pro199) in place of the residue that acts as donor for the second h-bond, impeding its formation. This chemical modification presumably contributes to destabilize the closed state in this BD. It is worth noting that in most crystal structures of ZMYND11 the ZA-loop is not resolved. In a notable exception (PDB 4N4G), authors proved that contacts with another crystallographic unit stabilize this flexible region, making it observable.^49^ This is consistent with our simulations, as we find the ZA-loop switching between the two states that are shown in Figure 3, making it difficult to capture its electron density. Importantly, this observation provides direct evidence of the possibility to modulate BD flexibility with macromolecular contacts, suggesting that similar interactions with other biological entities – *e.g*. DNA– could also lead to such conformational changes.

The other crystallographic evidence is a structure of PB1(6), which is also an atypical BD having an unusually short ZA-loop.^20^ In comparison with SMARCA2, a member of the same family, it shows a very similar opening despite having a low sequence identity (Figure 4c). The presence of a bulky threonine residue (Thr789) in place of the highly conserved aspartate may be one of the reasons why this BD is not stable in the closed state. We note that there are a few other BDs lacking this aspartate, and yet their crystal structures are stable in the closed state (see Supplementary Fig. 18). In these BDs the aspartate is replaced by residues like serine, alanine, or tryptophan, which represent drastic changes in terms of amino acid properties. Nonetheless, these modifications are accompanied by changes in the surrounding residues, leading to complementary interactions. This highlights that epistatic effects can compensate for the lack of the conserved aspartate, adapting local interactions to stabilize the closed state and retain BD function.

### Chemical shift predictions suggest a link between the novel state and a DNA-binding mode

Understanding the biological relevance of conformational states in a protein is crucial for the success of inhibitor design. However, this is not an easy task given the number of possible interactions that can occur in a cellular context. A recent study has demonstrated that SMARCA2 interacts with DNA through a basic surface patch, whose electrostatic contribution drive BD association to chromatin.^27^ Reported chemical shifts upon BD-DNA binding show significant perturbations in the region of the ZA-loop and the α_A_ helix, adjacent to where we observe the novel conformational change. This observation prompted us to compute the chemical shift of ^1^H, ^13^Cα and ^15^N atoms along the process, using the CAMSHIFT empirical predictor (Supplementary Fig. 19).^50^ Intriguingly, comparison of the predicted ^1^H shifts with the experimental values reveal a clear perturbation for Leu1412 (Fig. 5a), which is hydrogen bonded with the conserved aspartate (Supplementary Fig. 9a). This perturbation likely arise from contributions due to the conformational change along with the ones associated to DNA binding. In other words, these results suggest that SMARCA2 could undergo a population shift from the closed state to the open as a consequence of the interactions with DNA (Fig. 5b), modulating its flexibility and function in an allosteric manner. This hypothesis opens the door for future experiments to further describe the nature of the novel state and its relevance to DNA binding.

**Figure 5.**
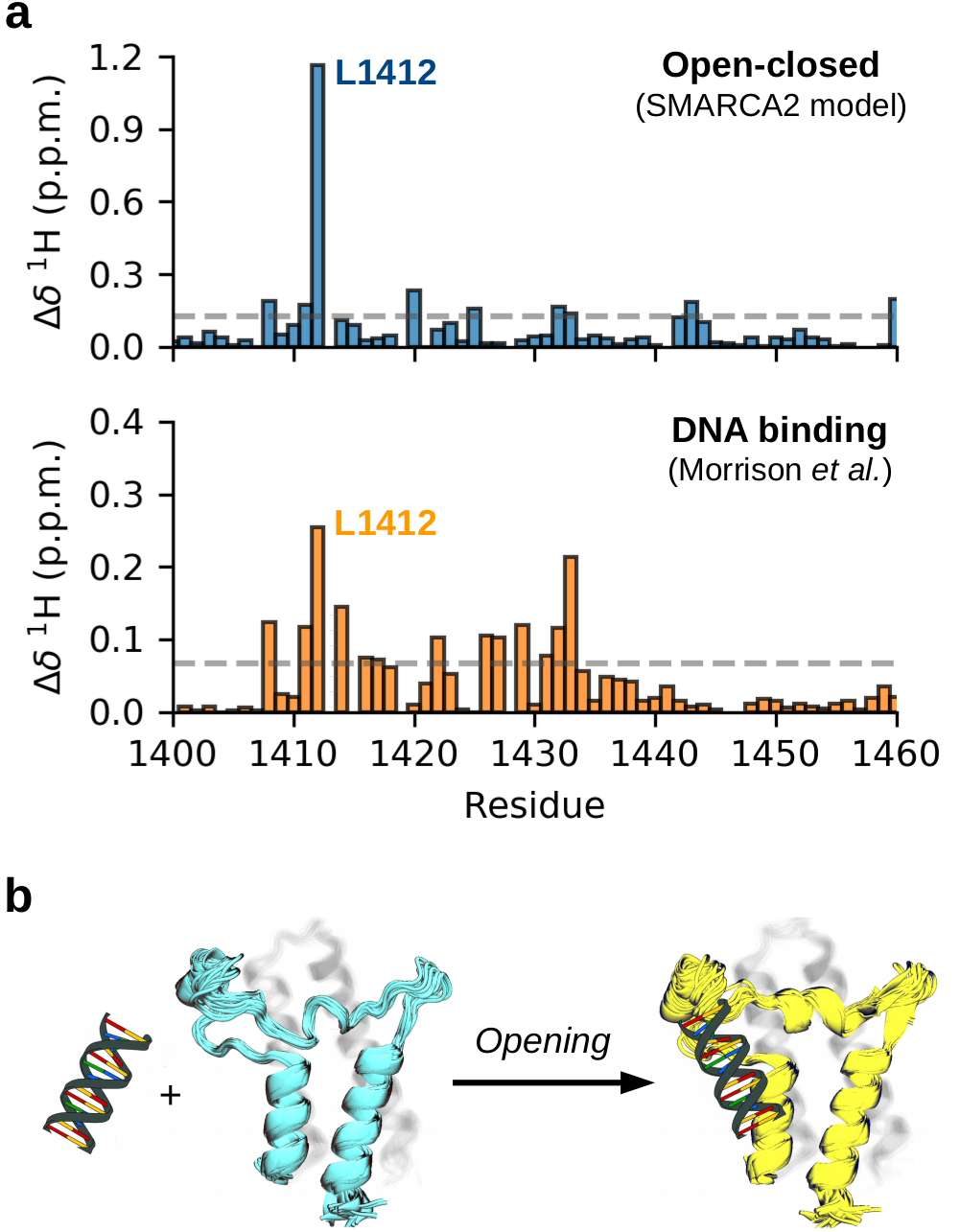
Chemical shift predictions of SMARCA2 along the conformational change correlate with a BD-DNA binding mode. (**a**) CAMSHIFT ^1^H predictions along the conformational change of SMARCA2 and experimental shifts upon its binding to DNA, obtained from Morrison *et al*. The numbering of residues is that of PDB 5DKC. (**b**) Hypothesis of a closed-to-open population shift as a consequence of BD interactions with DNA.

## DISCUSSION

In this work we have used extensive molecular dynamics simulations and Markov state modeling to characterize the dynamics and flexibility of bromodomains (BDs). Our results reveal a novel and ubiquitous state in which the ZA-loop displaces from the α_A_ helix, disrupting the ZA channel and opening a pocket beneath it. The conservation of this state across families, persisting along evolutionary pressures, suggests that it has a pivotal role in BD function and regulation.

Crystallographic structures projected onto a free energy map show an excellent agreement between the distribution of experimental structures and simulations. These results highlight that the conformational change is governed by a few conserved interactions among BDs, and reinforces previous studies that demonstrate the correspondence between ensembles of crystallographic structures and protein plasticity, both in solution and during enzyme catalysis.^51,52^ Furthermore, we report that two crystallographic structures, one of ZMYND11 and one of PB1(6), are stable in the open state. These structures directly support the existence of the novel state and reveal subtle changes of conserved residues that could help in the rationalization of disease related mutations. One of these changes is Pro199 in ZMYND11, which abrogates one of the key h-bonds that stabilize the closed state. This observation suggests that BDs could be engineered in the open state with a single site directed mutation, possibly switching between biologically active and inactive forms, with applications in target validation. Another change is Thr789 in PB1(6), which replaces the well-conserved aspartate, and whose mutation is in tumor suppressor genes in BDs of the same family (*e.g*. PB1(5) and the D705G variant).^53^

The novel state represents a potential target for future drug design campaigns, given the presence of cryptic pockets with allosteric effects. These pockets open in slightly different regions depending on the BD, offering alternative possibilities for interactions compared to those present in the acetyl-lysine pocket, being promising to overcome selectivity problems. Interestingly, one of the crystal structures of ZMYND11 (PDB 4N4I) shows a polyethylene glycol molecule bound beneath the predicted pocket, in the hydrophobic core of the protein, suggesting that the novel pockets may be indeed ligandable (see Supplementary Fig. 8d). This evidence further opens the question of whether different molecules can act as allosteric modulators in a cellular context. More generally, we envisage the possibility of controlling BD function via the design of specific inhibitors targeting these pockets.

Finally, we have proposed that the novel state possibly connects to a recently unveiled BD-DNA binding mode, which displays significant chemical shift perturbations in the opening region. We hypothesize that such biologically relevant interactions can shift the population between the states, stabilizing the open state to occlude the acetyl-lysine pocket and regulate BD function. Altogether, we believe that our results significantly contribute to the general understanding of BD structure, dynamics and function, providing a mechanism of potential autoregulation that deepens in BD biology and paves the way towards the design of future inhibitors with higher potency and selectivity.

## ACKNOWLEDGEMENTS

We acknowledge funding from the European Commission (ERC CoG 772230 “ScaleCell”), the Deutsche Forschungsgemeinschaft (CRC1114/C03, NO825/2-2), the MATH+ Berlin Mathematics Research Center (AA1×6, EF1×2), the Alexander von Humboldt Foundation (postdoctoral fellowship to S.O.), and Bayer AG (LSC project: “QuAnTaStics”). We thank Jacopo Negroni, Joseph Bluck, Michael Edmund Beck, Sikander Hayat, Jeremie Mortier, Holger Steuber and Nicolas Werbeck for feedback and discussions at different stages during this study. We also thank Emma A. Morrison and Catherine A. Musselman for sharing raw chemical shift data from reference 27.

