## Supporting Material for "Discovery of a hidden transient state in all bromodomain families"

### Supplementary Information
### Discovery of a hidden transient state in all bromodomain families

Lluís Raich,[a] Katharina Meier,[b] Judith Günther,[c] Clara D. Christ,[c] Frank Noé[a,d,*] and Simon Olsson[a,*]

[a]*Freie Universität Berlin, Department of Mathematics and Computer Science, Arnimallee 6, 14195 Berlin, Germany.* [b]*Bayer AG, Pharmaceuticals, R&D, Computational Molecular Design, 42096 Wuppertal,* Germany. [c]*Bayer AG, Pharmaceuticals, R&D, Computational Molecular Design, 13342 Berlin,* Germany. [d]*Rice University, Department of Chemistry, Houston, TX 77005, USA.*

#### Table of contents

1. Computational details and methodology
2. Markov state models of each bromodomain family
3. Analysis of free energy landscapes
4. Analysis of pocket volumes
5. Analysis of experimental structures
6. Nuclear magnetic resonance predictions
7. References

#### 1. Computational details and methodology

***Set up of the systems:*** bromodomain (BD) structures were obtained from the protein data bank (PDB), using the identifiers and amino acid sequences that are given in Supplementary Table 1. The structures of PHIP(2), BRD1, BAZ2B, TRIM28 and ZMYND11 were shortened in their N- or C-termini to reduce system size. The structure of ZMYND11, from mice and with part of the ZA-loop unresolved, was completed using the Swiss-Model server for homology modeling,[1] modifying the only residue that differs between humans and mice (Ser183Asn). All structures were cleaned of cosolvent molecules and drugs while retaining crystallographic waters. Hydrogen atoms and additional solvent molecules were added using the tleap module of Ambertools[2] (v 19.0), with a distance of 1 nm from any atom of the protein and the simulation box. Counterions were added to neutralize the total charge of the systems and set a 0.15 M NaCl salt concentration. We used the ff14SB[3] force field together with the TIP3P[4] water model to describe the solute and the solvent, respectively. This force field has been previously used to successfully compute protein-ligand affinities in BRD4(1), with errors below 3 kcal/mol.[5]

***Simulation details:*** molecular dynamics simulations were carried out with the OpenMM[6] software (v 7.4.0). A Langevin integrator with a 1 ps friction coefficient and a Monte Carlo barostat were used to maintain the temperature and pressure of the systems at 300 K and 1 bar. Long-range electrostatics were computed using the Particle-Mesh Ewald scheme, and van der Waals interactions were truncated with a 1 nm cutoff. A hydrogen mass repartition with a factor of 2 –including both solute and solvent atoms– was used to increase the integration time step up to 4 fs, constraining the lengths of all bonds. Multiple 1 µs trajectories were ran for each BD, with an initial minimization and 5 additional nanoseconds of equilibration for each trajectory. Frames were saved every 100 ps, including all atoms of the system. As an initial sampling strategy, for each BD we ran 4 trajectories of 250 ns at a higher temperature, two at 350 K and two at 375 K. From these simulations we selected snapshots based on $C_\alpha$ RMSD (root-mean-square deviation; with the crystallographic structure as reference) to run the production simulations at 300 K. The structure of ZMYND11 was not forced with this temperature jump since it was already in the open state and part of the ZA-loop was modeled. An adaptive sampling strategy was adopted afterwards, estimating preliminary Markov state models and selecting snapshots focused on the region of interest.

Trajectories were superimposed onto the crystallographic structure based on $C_\alpha$ RMSD using the tleap module, and all solvent atoms and ions were stripped for the subsequent analyses.

| Family | I | II | III | IV |
|---|---|---|---|---|
| BD name | CECR2 | BRD4(1) | PHIP(2) | BRD1 |
| PDB id | 3NXB | 5ULA | 3MB3 | 3RCW |
| N-terminus | Asp438 | Asn44 | Ala1320 | Thr565 |
| C-terminus | Lys537 | Asn162 | Leu1422 | Val670 |
| Trajectories (μs) | 40 | 64 | 60 | 40 |

| Family | V | VI | VII | VIII |
|---|---|---|---|---|
| BD name | BAZ2B | TRIM28 | ZMYND11 | SMARCA2 |
| PDB id | 3G0L | 2RO1 | 4N4G | 5DKC |
| N-terminus | Leu1870 | Ser697 | Gln156 | Pro1381 |
| C-terminus | Lys1970 | Phe800 | Gln256 | Lys1490 |
| Trajectories (μs) | 40 | 40 | 48 | 60 |

**Supplementary Table 1.** Bromodomains, protein data bank identifiers and amino acid sequences used in this work, together with the total number of 1 μs trajectories.

***Markov modeling:*** Markov state models (MSM) were built for each individual BD using the PyEMMA[7] software (v 2.5.7), following well-established workflows and protocols.[8] First, we modeled BRD4(1) taking into account residues Leu67 to Arg113 and using different sets of features, including 41 backbone h-bond distances (between heteroatoms) formed along each trajectory, 1081 $C_\alpha$ pairwise distances, 141 $C_\alpha$ XYZ coordinates and 188 sin/cos backbone torsion angles. Trajectories 17 and 25 were filtered from the total since they are involved in a slow process that is related with the loss of a conserved π-stacking interaction (Tyr97-Tyr139). We used a time-lagged independent component analysis (TICA[9,10]) to reduce the dimensionality of each set of features, using a lag time of 10 ns and taking 6 to 14 components depending on the input features (80 to 95% of cumulative variance). The reduced spaces were clustered using the k-means algorithm with 100 centers, and Bayesian MSMs were built selecting the lag times according to the convergence of the implied timescales (0.1 μs). The models were coarse-grained to two states using Perron cluster cluster analysis (PCCA++) and validated with a Chapman-Kolmogorov (CK) test. Errors in populations, mean first passage times (MFPT) and CK tests were computed using a bootstrapping strategy without replacement (100 rounds for populations and MFPTs; 10 for CK tests), taking 80% of the initial trajectories and fitting a new MSM using the same feature map, clustering and metastable discretization than the MSM with the total set of trajectories. All these models showed a perfect agreement for the prediction of thermodynamic and kinetic properties within statistical uncertainties (Supplementary Figure 1), highlighting the robustness of the results independently of the input features, the number of time-lagged independent components (TICs), clustering and coarse-grain discretization. We also reduced the model of backbone h-bonds from 41 input distances to the two most relevant for describing the first TIC (Phe83-Met107 and Gln84-Gly108 backbone h-bonds), finding that this simplified model captures all the kinetic information of the system along the process. Therefore, we decided to use this intuitive featurization to describe the kinetics of all BDs, aiming to facilitate the interpretation of the models and their mutual comparison.

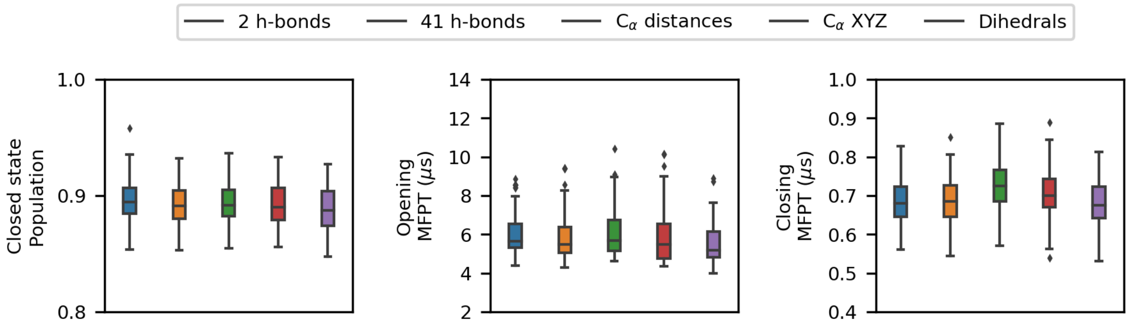


**Supplementary Figure 1. Consistency of thermodynamic and kinetic predictions for BRD4(1) with different features.** Boxplots of a bootstrapped distribution of the closed state population and opening/closing mean first passage time (MFPT) for the two key backbone h-bonds (2 TICs), 41 backbone h-bonds (6 TICs), 1081 $C_\alpha$ distances (13 TICs), 141 $C_\alpha$ XYZ coordinates (14 TICs) and 188 sin/cos backbone dihedral pairs (10 TICs).

We obtained a general map of TICs using the two key backbone h-bonds and all the trajectories of each BD, defining a lag time of 10 ns and taking both dimensions. The first component represents the breaking of the two h-bonds and is described by the linear combination 2.16·(h-bond 1)+0.20·(h-bond 2), while the second represents the formation of h-bond 1 together with the breaking of h-bond 2 and is described by the linear combination -5.09·(h-bond 1)+5.69·(h-bond 2). This general TIC space was used to build MSMs for each BD. To minimize projection errors in this low-dimensional space, we filtered trajectories that were not reversibly connected in a higher dimensional space made by $C_\alpha$ pairwise distances for each BD, as well as trajectories involved in other slow processes that can not be properly resolved in the reduced space. A pair of metastable states are reversible connected if empirical transitions in both directions have been observed at least once. In total, trajectories 11, 13 and 20 were filtered for CECR2; 17 and 25 for BRD4(1); 1, 14-18, 20, 35, 44, 47, 49, 52, 53 and 57 for PHIP(2); 7, 8, 12, 26, 31, 32, 34, 35 and 39 for BRD1; 9-11, 28, 30-40 for BAZ2B; 37 for TRIM28; 28, 30, 31, 33, 39 and 47 for ZMYND11 and 32, 46, 47, 49, 52 and 53 for SMARCA2. These trajectories involve a diverse set of slow conformational changes, such as the loss of a conserved π-stacking interaction (*e.g.* Tyr471-Tyr513 in CECR2), the perturbation of a conserved hydrophobic residue that is next to the conserved aspartate (*e.g.* Met479 in CECR2), or transitions between different open state conformations of the ZA-loop for BDs that display a high disorder in this region (*e.g.* PHIP(2), BRD1 and BAZ2B). The filtered trajectories were mapped onto the general TIC space and MSMs were built following the same protocols given above for BRD4(1), including an additional convergence test plotting state populations with respect to the lag time. Structural details, kinetic parameters and convergence tests for each of the models are given in Supplementary Figures 2-9. Populations and MFPTs of the bootstrapping distributions are given in Supplementary Tables 2 and 3. Representative structures were selected by taking the most populated cluster of each metastable state, drawing 1000 random snapshots assigned to it. The conformational complexity of TRIM28 prevented us from building a reliable MSM with the acquired simulation data, and thus we only provide evidence of metastability for the open state.

***Analysis of crystal structures:*** bromodomain structures of the Pfam[11] database (PF00439) were downloaded from the PDB and superimposed with Theseus[12] (v 3.3.0), using Muscle (v 3.8.1) for the alignment of sequences. This superimposition and alignment was used to identify the key residues of each BD that are involved in the two backbone h-bonds, and we subsequently computed their distances using MDtraj. Due to ambiguity in the sequence alignment, we manually inspected the structures of all outliers (half of the standard deviation above the median), and in the case of a misassignment we recalculated h-bond distances updating the key residues. In ZMYND11 the identification of such residues is not clear given that in the experimental structures the ZA-loop is either completely unresolved (*e.g.* PDB 4N4I) or partially resolved and in a very open state (PDB 4N4G). The only requirement for the h-bond 1 acceptor residue is that it has to be non-polar and preferably hydrophobic, since it should be bound in the hydrophobic core of the protein when closed. For this reason, we selected Gly180 to act as acceptor for h-bond 1, as it is the only neutral residue in the central part of the ZA-loop that is resolved

(PDB 4N4G). Supplementary Figure 17 shows the projection of all structures onto the raw free energy landscape of each BD along the two key h-bonds. Supplementary Figure 18 shows four experimental structures of BDs lacking the conserved aspartate, with different amino acids (Ala, Ser, Trp and Thr) at the critical position. A table with all PDB identifiers, residues and distances is provided as online material.

***Additional details:*** the visual inspection of structures and the production of figures were done with VMD[13] (v 1.9.3). MDtraj[14] (v 1.9.3) was used for computing structural parameters, including h-bond analyses and solvent accessible surface areas (SASA). Plumed[15] (v 2.5.3) was used for the prediction of chemical shifts within the CAMSHIFT[16] model. NGLview[17] (v 2.7.1) was used to visualize structures during preliminary analyses. All plots were done with Matplotlib (v 3.1.1) and the seaborn library (v 0.9.0) was used for the distributions –with a Gaussian kernel density estimation– and boxplots.

MDpocket[18] (v 3.0) was used to detect small molecule binding sites and to calculate pocket volumes (Supplementary Figure 15). Acetyl-lysine pockets were manually selected from each closed state using frequency maps at isovalues of 0.5, corresponding to at least 50% of pocket opening in all frames. For PHIP(2) the acetyl-lysine pocket was selected at isovalues of 0.25 given its low opening frequency, discarding parts that extend towards the connection between the ZA-loop and the $\alpha_A$ helix. For ZMYND11 we selected the principal cryptic pocket of the open state since the closed state is never fully formed and thus the acetyl-lysine pocket is not defined (see Supplementary Figure 15b). The shrinking of pockets was estimated computing their volumes with the structures of both closed and open states, discarding null volumes that result as a consequence of pocket parameters (minimum of 35 alpha spheres and 3 Å radius). Note that the evaluation of acetyl-lysine pocket volumes with the structures of the open state gives an estimate of their occlusion as a consequence of the conformational change.

The sequence alignment in Figure 1b was done using the STAMP[19] algorithm implemented in VMD, which takes advantage of structural information. First, we aligned the ZA-loop segments of all minimized structures, resulting an alignment with several gaps given the unusual conformations of TRIM28 and ZMYND11. In a second step, we refined the sequence of ZMYND11 manually, filling the gaps of the backbone h-bond acceptors with the nearest residues in the alignment (Gly180 and Lys181). Finally, we realigned the entire sequence of TRIM28 using ClustalX[20] (v 2.1) and removed gaps in the entire alignment that lead to single amino acids disconnected.

#### 2. Markov state models of each bromodomain family

In this section we provide a summary of the Markov state models of each BD family (Supplementary figures 2 to 9). All figures are organized identically with 5 different subsections, apart from TRIM28 and ZMYND11. The first subsection (a) shows the general BD fold and a close view of the key region for the two metastable states, closed and open, colored in cyan and yellow respectively. The second and the third (b and c) show structural parameters, free energy profiles and mean-first passage times that characterize the conformational change. The fourth (d) shows a superposition of the predicted closed states and their pocket frequency maps with experimental structures of BDs complexed with drugs. Finally, the fifth subsection (e) shows convergence tests for the MSMs, including the Chapman-Kolmogorov test and time convergence of populations and timescales. TRIM28 and ZMYND11 lack convergence tests given that we do not use their MSMs to report populations nor timescales. Furthermore, TRIM28 lacks the section of pocket frequency maps as it does not display noticeable pockets in the region of interest. Instead of this, we provide the evolution of three selected trajectories that show clear metastability in the open state.

We note that for CECR2, BRD4(1) and SMARCA2 systems we generally see a clear two-state behavior, with very stable structures and few alternative states, resulting in well converged models. On the other hand, PHIP(2), BRD1 and BAZ2B systems have more complex conformational landscapes, and thus the two-state approximation and the simplified featurization may affect the robustness of the reported rates and state populations. This can be observed in the Chapman-Kolmogorov tests of these systems (Supplementary Figures 4-6 panel e), which show large confidence intervals and a moderate agreement between the predicted and estimated decay of probability densities. Nevertheless, all models clearly resolve the metastability of the closed and open states of the ZA-loop, confirming its kinetic relevance.

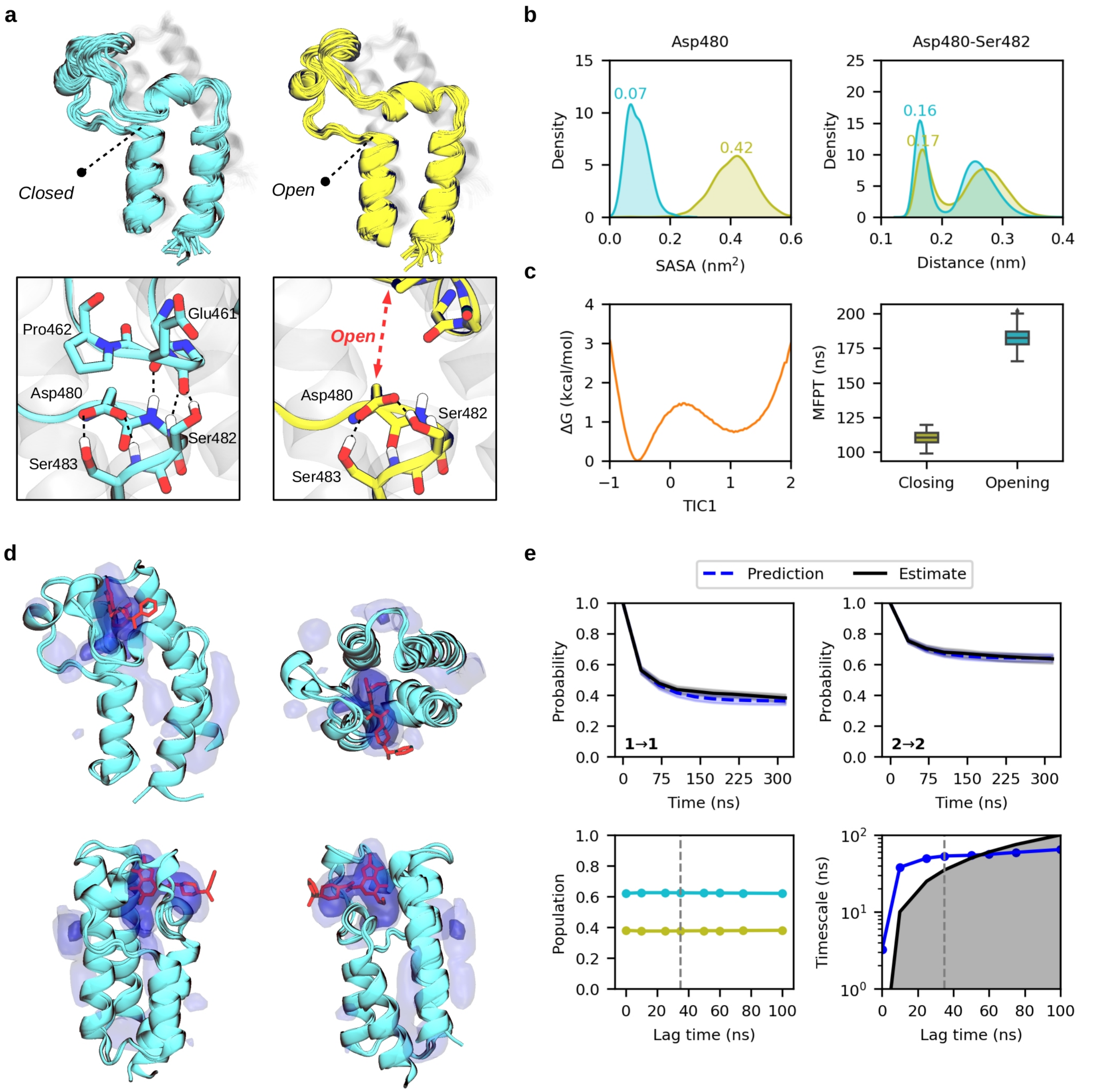


**Supplementary Figure 2. The novel conformational state in CECR2 of family I.** (**a**) Ensemble of structures of the "closed" crystal-like state (cyan) and the "open" novel state (yellow). A close view of the opening region is represented below, highlighting Asp480. (**b**) Distribution of the Asp480 solvent accessible surface area (SASA) and closest distance between Asp480-Ser482 side chains for the two metastable states. (**c**) Reweighted free energy profile along the second component (TIC1) and boxplot with the opening and closing mean first passage times. (**d**) Pocket frequency maps of the closed state superposed with an X-ray structure (PDB 5V84) in complex with a drug molecule shown in red. Isosurfaces are given at 0.25 (light) and 0.50 (intense) values. (**e**) Chapman-Kolmogorov test for the two states (1 open, 2 closed), and convergence of populations and timescales with respect to the lag time. Vertical dashed lines indicate the lag time used to build the MSM (35 ns).

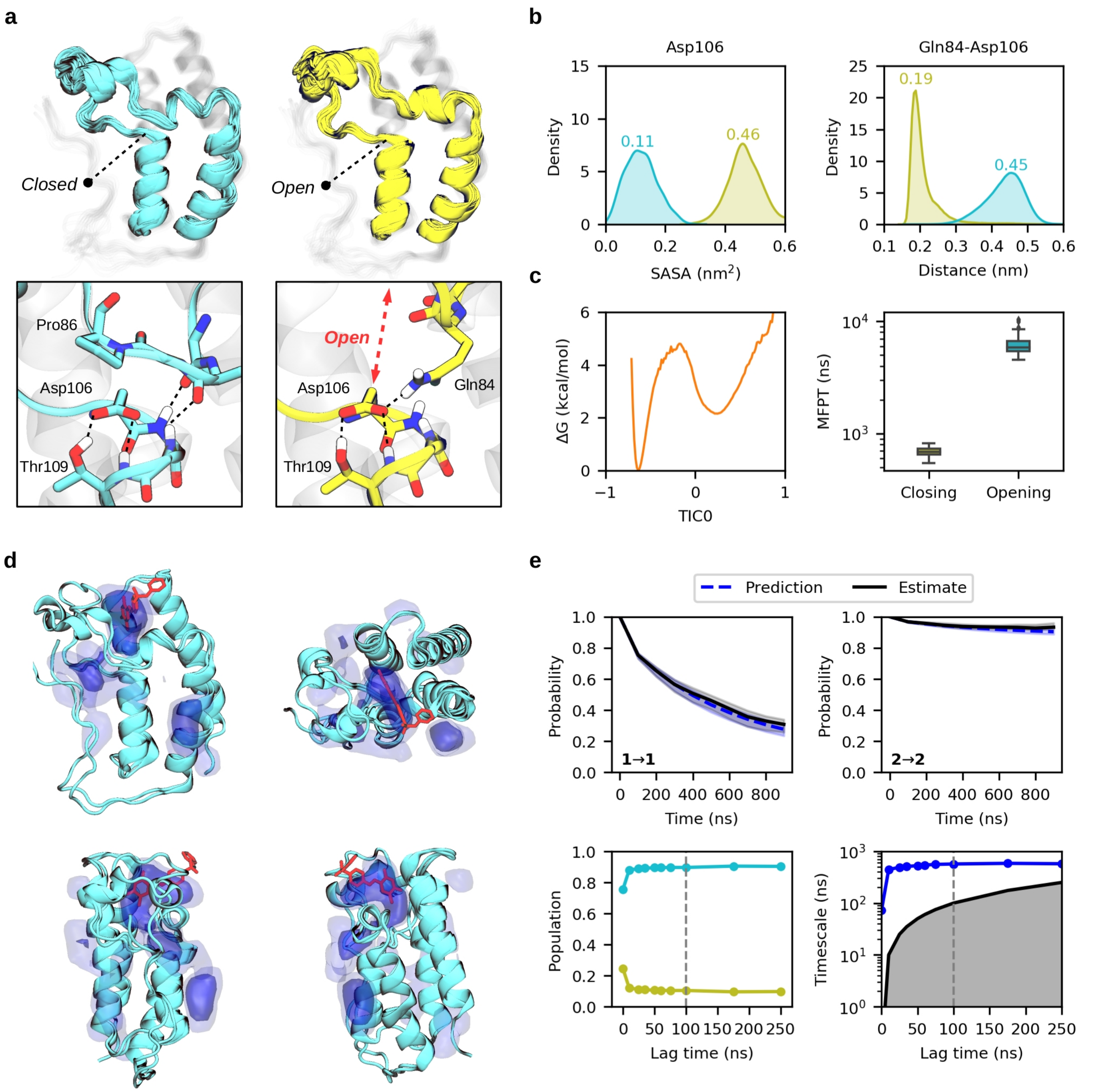


**Supplementary Figure 3. The novel conformational state in BRD4(1) of family II.** (**a**) Ensemble of structures of the "closed" crystal-like state (cyan) and the "open" novel state (yellow). A close view of the opening region is represented below, highlighting Asp106. (**b**) Distribution of the Asp106 solvent accessible surface area (SASA) and closest distance between Gln86-Asp106 side chains for the two metastable states. (**c**) Reweighted free energy profile along the first component (TIC0) and boxplot with the opening and closing mean first passage times. (**d**) Pocket frequency maps of the closed state superposed with an X-ray structure (PDB 4NUD) in complex with a drug molecule shown in red. Isosurfaces are given at 0.25 (light) and 0.50 (intense) values. (**e**) Chapman-Kolmogorov test for the two states (1 open, 2 closed), and convergence of populations and timescales with respect to the lag time. Vertical dashed lines indicate the lag time used to build the MSM (100 ns).

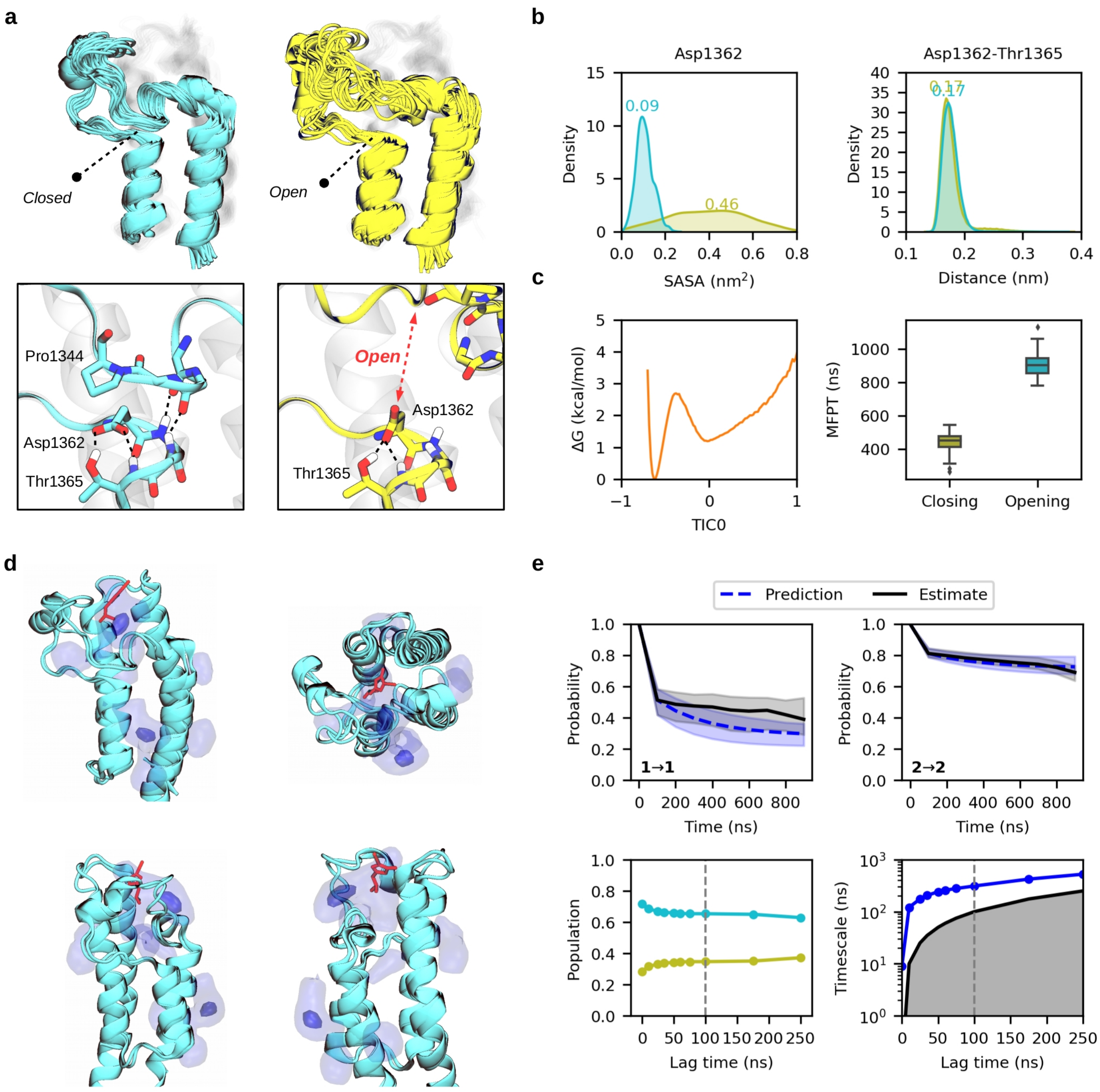


**Supplementary Figure 4. The novel conformational state in PHIP(2) of family III.** (**a**) Ensemble of structures of the "closed" crystal-like state (cyan) and the "open" novel state (yellow). A close view of the opening region is represented below, highlighting Asp1362. (**b**) Distribution of the Asp1362 solvent accessible surface area (SASA) and closest distance between Asp1362-Thr1365 side chains for the two metastable states. (**c**) Reweighted free energy profile along the first component (TIC0) and boxplot with the opening and closing mean first passage times. (**d**) Pocket frequency maps of the closed state superposed with an X-ray structure (PDB 5ENF) in complex with a molecular fragment shown in red. Isosurfaces are given at 0.25 (light) and 0.50 (intense) values. (**e**) Chapman-Kolmogorov test for the two states (1 open, 2 closed), and convergence of populations and timescales with respect to the lag time. Vertical dashed lines indicate the lag time used to build the MSM (100 ns).

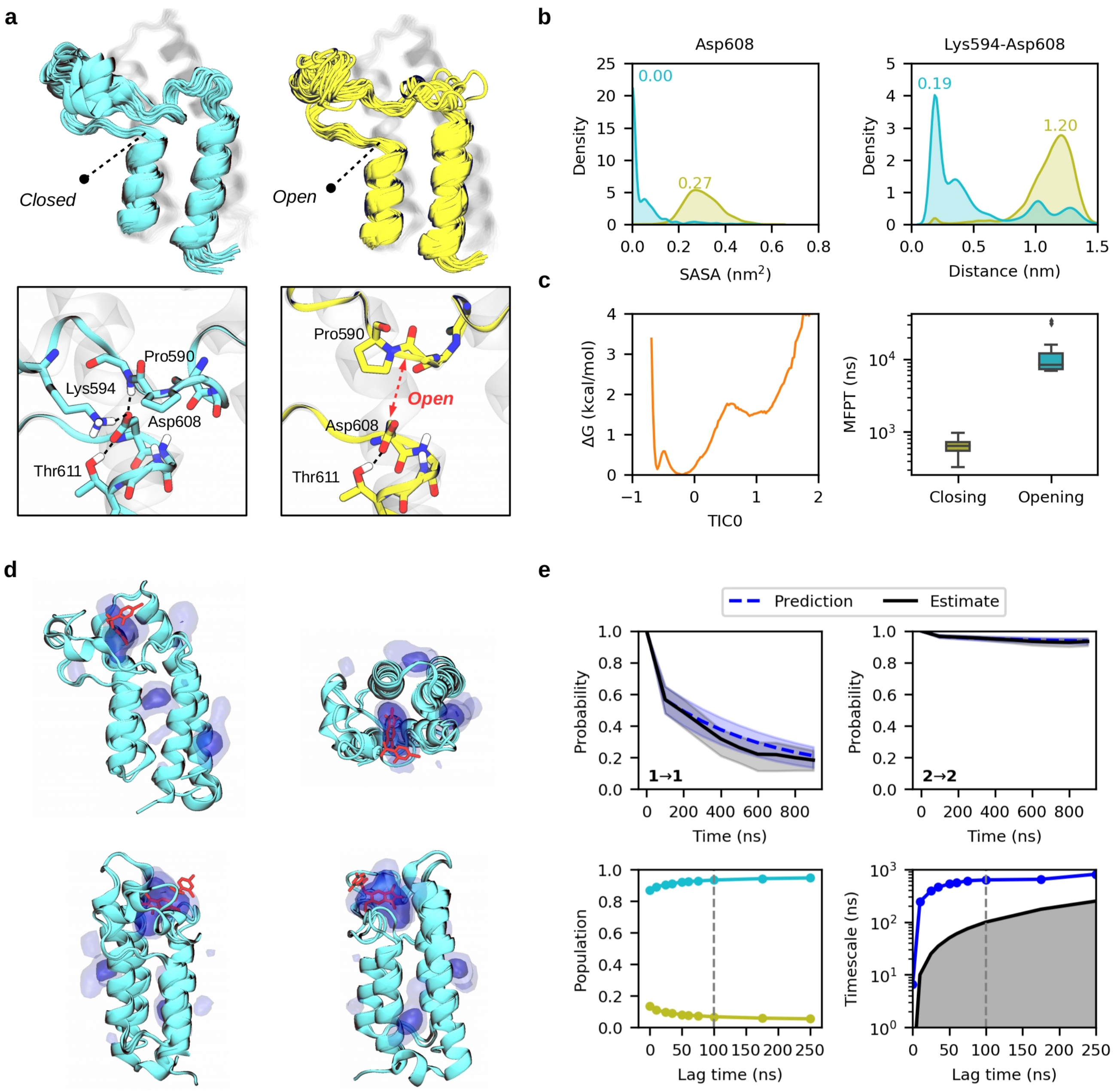


**Supplementary Figure 5. The novel conformational state in BRD1 of family IV.** (**a**) Ensemble of structures of the "closed" crystal-like state (cyan) and the "open" novel state (yellow). A close view of the opening region is represented below, highlighting Asp608. (**b**) Distribution of the Asp608 solvent accessible surface area (SASA) and closest distance between Lys594-Asp608 side chains for the two metastable states. (**c**) Reweighted free energy profile along the first component (TIC0) and boxplot with the opening and closing mean first passage times. (**d**) Pocket frequency maps of the closed state superposed with an X-ray structure (PDB 5FG6) in complex with a drug molecule shown in red. Isosurfaces are given at 0.25 (light) and 0.50 (intense) values. (**e**) Chapman-Kolmogorov test for the two states (1 open, 2 closed), and convergence of populations and timescales with respect to the lag time. Vertical dashed lines indicate the lag time used to build the MSM (100 ns).

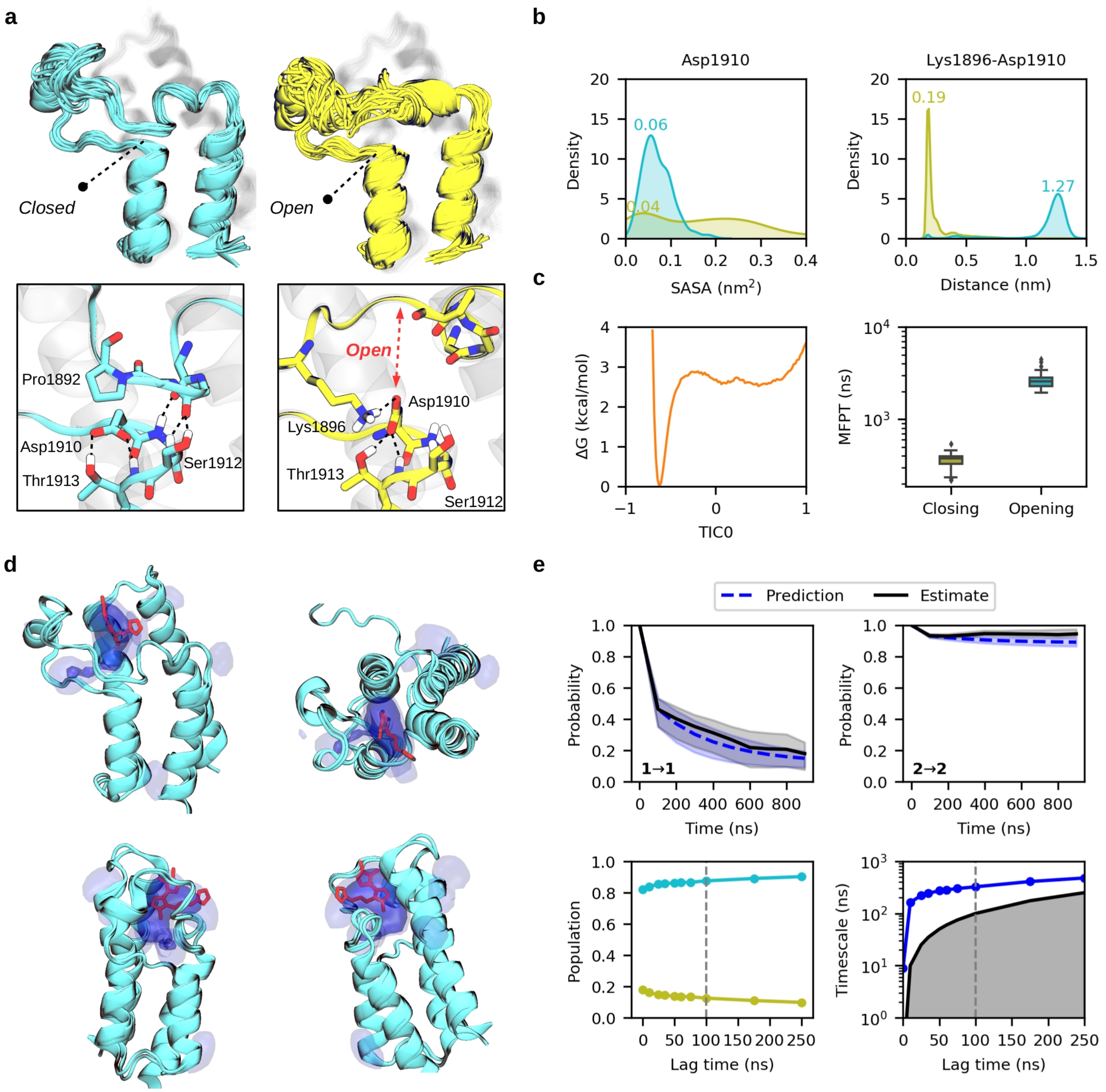


**Supplementary Figure 6. The novel conformational state in BAZ2B of family V.** (**a**) Ensemble of structures of the "closed" crystal-like state (cyan) and the "open" novel state (yellow). A close view of the opening region is represented below, highlighting Asp1910. (**b**) Distribution of the Asp1910 solvent accessible surface area (SASA) and closest distance between Lys1896-Asp1910 side chains for the two metastable states. (**c**) Reweighted free energy profile along the first component (TIC0) and boxplot with the opening and closing mean first passage times. (**d**) Pocket frequency maps of the closed state superposed with an X-ray structure (PDB 3Q2F) in complex with a drug molecule shown in red. Isosurfaces are given at 0.25 (light) and 0.50 (intense) values. (**e**) Chapman-Kolmogorov test for the two states (1 open, 2 closed), and convergence of populations and timescales with respect to the lag time. Vertical dashed lines indicate the lag time used to build the MSM (100 ns).

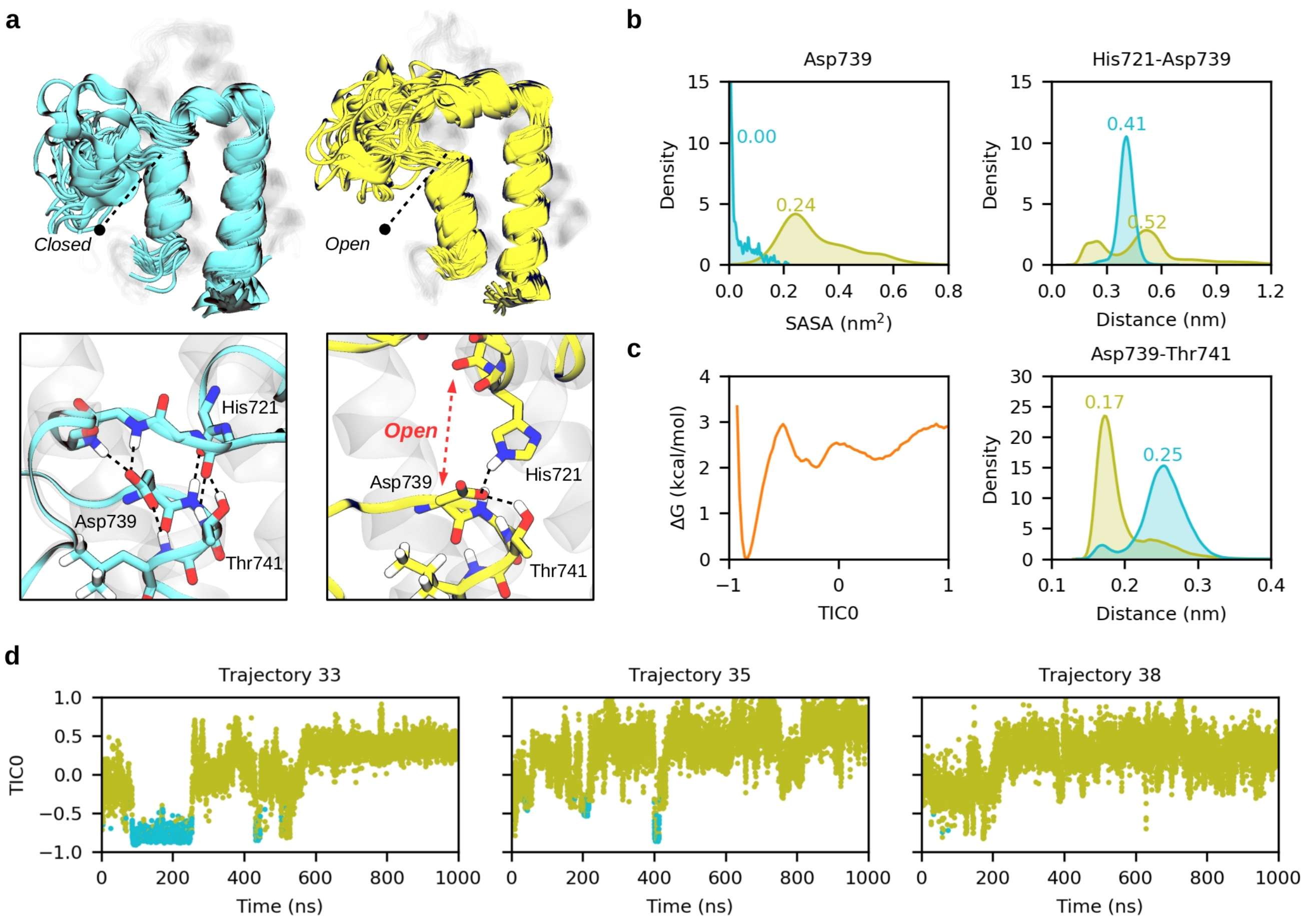


**Supplementary Figure 7. The novel conformational state in TRIM28 of family VI.** (**a**) Ensemble of structures of the "closed" crystal-like state (cyan) and the "open" novel state (yellow). A close view of the opening region is represented below, highlighting Asp739. (**b**) Distribution of the Asp739 solvent accessible surface area (SASA) and closest distance between His721-Asp739 side chains for the two metastable states. (**c**) Raw free energy profile along the first component (TIC0) and distribution of the Asp739-Thr741 closest distance for the two metastable states. (**d**) Evolution of TIC0 along three independent 1 μs trajectories, showing clear metastability between the two states.

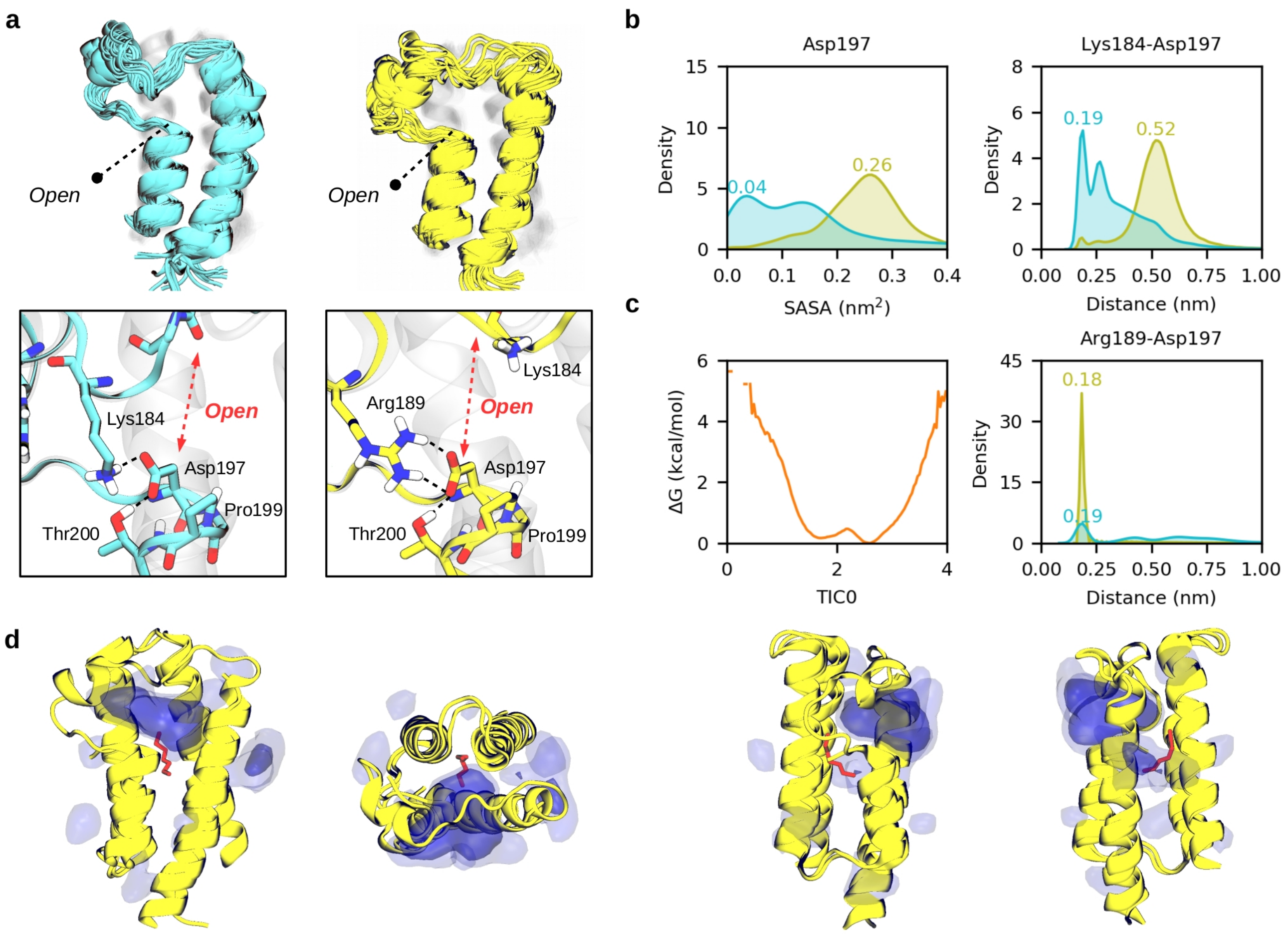


**Supplementary Figure 8. The novel conformational state in ZMYND11 of family VII.** (**a**) Ensemble of structures of the “semiclosed” state (cyan) and the “open” crystal-like state (yellow). A close view of the opening region is represented below, highlighting Asp197. (**b**) Distribution of the Asp197 solvent accessible surface area (SASA) and closest distance between Lys184-Asp197 side chains for the two metastable states. (**c**) Raw free energy profile along the first component (TIC0) and distribution of the Arg189-Asp197 closest distance for the two metastable states. (**d**) Pocket frequency maps of the open state superposed with an X-ray structure (PDB 4N4I) in complex with a polyethylene glycol molecule shown in red. Isosurfaces are given at 0.25 (light) and 0.50 (intense) values.

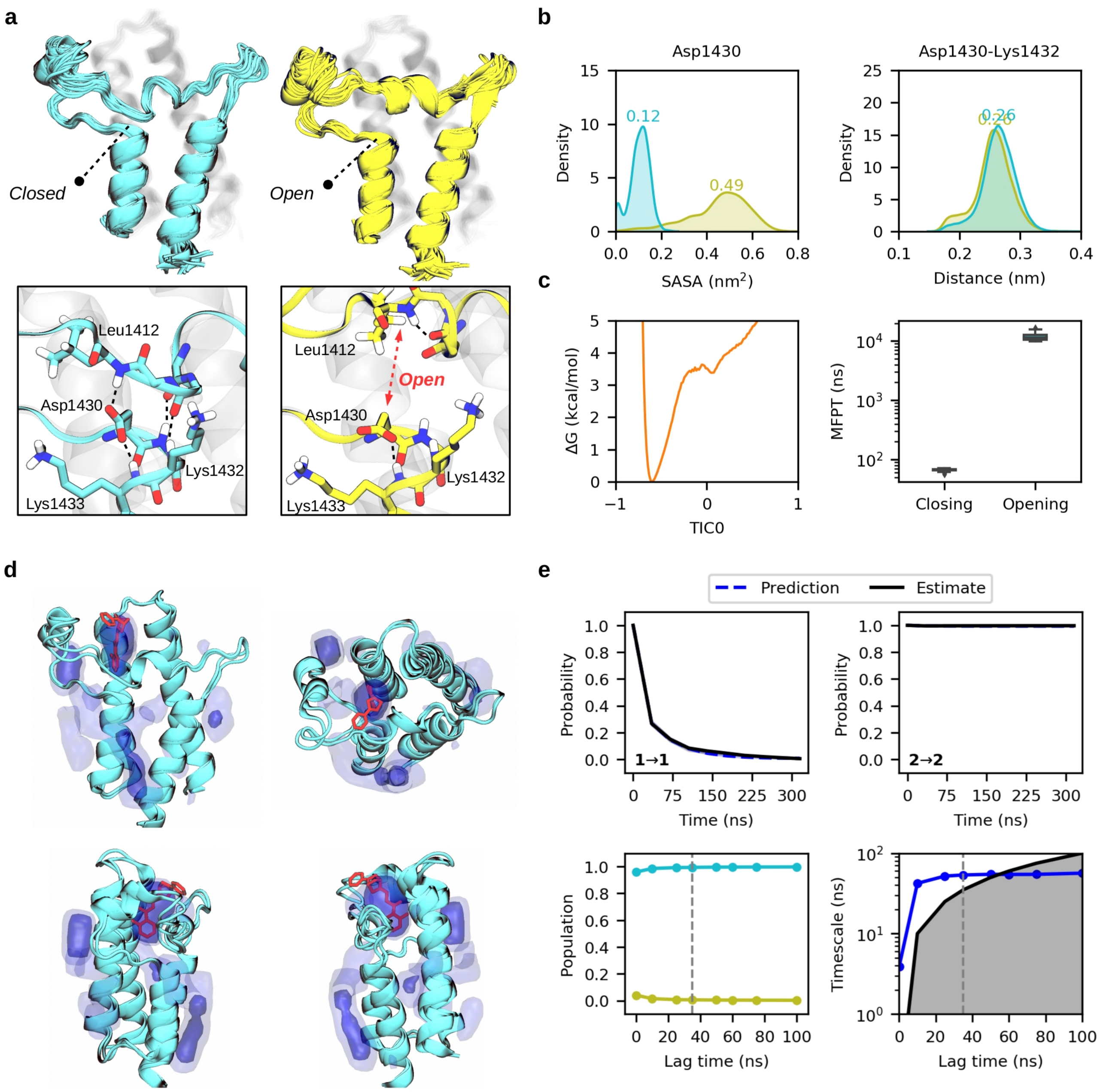


**Supplementary Figure 9. The novel conformational state in SMARCA2 of family VIII.** (**a**) Ensemble of structures of the "closed" crystal-like state (cyan) and the "open" novel state (yellow). A close view of the opening region is represented below, highlighting Asp1430. (**b**) Distribution of the Asp1430 solvent accessible surface area (SASA) and closest distance between Asp1430 -Lys1432 side chains for the two metastable states. (**c**) Reweighted free energy profile along the first component (TIC0) and boxplot with the opening and closing mean first passage times. (**d**) Pocket frequency maps of the closed state superposed with an X-ray structure (PDB 5DKC) in complex with a drug molecule shown in red. Isosurfaces are given at 0.25 (light) and 0.50 (intense) values. (**e**) Chapman-Kolmogorov test for the two states (1 open, 2 closed), and convergence of populations and timescales with respect to the lag time. Vertical dashed lines indicate the lag time used to build the MSM (35 ns).

| | CECR2 | | BRD4(1) | | PHIP(2) | |
|---|---|---|---|---|---|---|
| | Open | Closed | Open | Closed | Open | Closed |
| Samples | 100 | 100 | 100 | 100 | 100 | 100 |
| Mean | 0.38 | 0.62 | 0.10 | 0.90 | 0.34 | 0.66 |
| SD | 0.01 | 0.01 | 0.02 | 0.02 | 0.04 | 0.04 |
| Min | 0.34 | 0.59 | 0.06 | 0.85 | 0.21 | 0.59 |
| 25% | 0.37 | 0.61 | 0.09 | 0.88 | 0.32 | 0.63 |
| Median | 0.38 | 0.62 | 0.10 | 0.90 | 0.34 | 0.66 |
| 75% | 0.39 | 0.63 | 0.12 | 0.91 | 0.37 | 0.68 |
| Max | 0.41 | 0.66 | 0.15 | 0.94 | 0.41 | 0.79 |

| | BRD1 | | BAZ2B | | SMARCA2 | |
|---|---|---|---|---|---|---|
| | Open | Closed | Open | Closed | Open | Closed |
| Samples | 100 | 100 | 100 | 100 | 100 | 100 |
| Mean | 0.07 | 0.93 | 0.13 | 0.87 | 0.01 | 0.99 |
| SD | 0.02 | 0.02 | 0.02 | 0.02 | 0.00 | 0.00 |
| Min | 0.01 | 0.87 | 0.07 | 0.84 | 0.00 | 0.99 |
| 25% | 0.05 | 0.92 | 0.12 | 0.86 | 0.01 | 0.99 |
| Median | 0.07 | 0.93 | 0.13 | 0.87 | 0.01 | 0.99 |
| 75% | 0.08 | 0.95 | 0.14 | 0.88 | 0.01 | 0.99 |
| Max | 0.13 | 0.99 | 0.16 | 0.93 | 0.01 | 1.00 |

**Supplementary Table 2.** Bootstrapping distribution of populations for each BD, including the number of samples, mean and standard deviation (SD), minimum and maximum values of the distribution, median and the 25-75th percentiles.

| | CECR2 | | BRD4(1) | | PHIP(2) | |
|---|---|---|---|---|---|---|
| | Closing | Opening | Closing | Opening | Closing | Opening |
| Samples | 100 | 100 | 100 | 100 | 100 | 100 |
| Mean | 110 | 183 | 690 | 6129 | 441 | 907 |
| SD | 4 | 7 | 61 | 1233 | 51 | 70 |
| Min | 99 | 166 | 545 | 4564 | 262 | 781 |
| 25% | 107 | 178 | 651 | 5408 | 410 | 856 |
| Median | 110 | 182 | 688 | 5864 | 449 | 903 |
| 75% | 114 | 187 | 732 | 6667 | 477 | 945 |
| Max | 120 | 202 | 818 | 10418 | 544 | 1131 |

| | BRD1 | | BAZ2B | | SMARCA2 | |
|---|---|---|---|---|---|---|
| | Closing | Opening | Closing | Opening | Closing | Opening |
| Samples | 100 | 100 | 100 | 100 | 100 | 100 |
| Mean | 645 | 9936 | 364 | 2620 | 67 | 11784 |
| SD | 117 | 4096 | 57 | 455 | 3 | 1522 |
| Min | 327 | 6998 | 218 | 1930 | 56 | 9852 |
| 25% | 551 | 7312 | 329 | 2284 | 66 | 10420 |
| Median | 643 | 8344 | 377 | 2585 | 68 | 11523 |
| 75% | 721 | 12107 | 395 | 2809 | 69 | 12609 |
| Max | 971 | 33633 | 537 | 4487 | 71 | 16571 |

**Supplementary Table 3.** Bootstrapping distribution of mean-first passage times (ns) for each BD, including the number of samples, mean and standard deviation (SD), minimum and maximum values of the distribution, median and the 25-75th percentiles.

##### 3. Analysis of free energy landscapes

In this section we analyze the empirical free energy landscapes of each key h-bond and a contact distance that qualitatively describes the ZA-loop opening (Supplementary Figures 10 and 11). This contact distance is defined as the minimum distance between a segment of backbone heavy atoms of the ZA-loop and all atoms of the conserved aspartate. The segment extends from the ZA channel to a conserved hydrophobic residue that wraps the ZA-loop with the $\alpha_B$ helix (Glu461-Ile475 in CECR2; analogous residues in the other BDs).

The free energy landscapes of h-bond 1 and the ZA-loop opening (Supplementary Figure 10) display four interesting points to note. First, all maps show basins in the upper right quadrant (breaking of the h-bond and ZA-loop opening) except for CECR2, whose basin is more centered in the upper left quadrant (opening of the ZA-loop without breaking the h-bond). Second, the basins of ZMYND11 are found at values above 1 nm for the h-bond axis, highlighting that it does not form. Third, BRD1 has a low energy region that encompasses an equilibrium between formed and broken h-bond in the closed state. Fourth, most of the BDs substantially explore the upper left quadrant without showing notable minima, indicating opening events that rapidly close.

Regarding the third point, Supplementary Figure 12 shows detailed structures along the low energy region in BRD1, related with a slight displacement of the ZA-loop towards the $\alpha_Z$ helix. This displacement implies the breaking of the key h-bonds, which are compensated by additional h-bonds that backbone amides of the ZA-loop establish with the conserved aspartate. Interestingly, we found similar interactions in CECR2, PHIP(2), and BAZ2B (lower right quadrant in either Supplementary Figures 10 or 11), resembling the type of interactions that are present in the experimental structures of TRIM28 and all Family VIII members.

Regarding the fourth point, Supplementary Figure 13 shows trajectories of each BD were sporadic openings occur, characterized by narrow and dark peaks. These fast openings are common in all BDs and are generally not metastable. BRD1 is an exception, as it displays two open states that are metastable, with clear basins in the free energy landscape (one at 0.5 nm of h-bond 1 and the other at 1.0 nm; see Supplementary Figure 14). The state at 0.5 nm relaxes within a timescale that is faster than the resolution of the MSM, while the state at 1.0 nm is the slow process resolved by the model. The main difference between the two states is the orientation of the ZA-loop, which points outwards for the fast process and inwards for the slow, occluding the acetyl-lysine pocket.

Supplementary Table 4 reflects some of the points highlighted above, showing for CECR2 a substantial percentage of h-bond 1 in the open state, and for BRD1 a low percentage of the two key h-bonds in the closed state.

| | CECR2 | | BRD4(1) | | PHIP(2) | | BRD1 | |
|---|---|---|---|---|---|---|---|---|
| | h-bond 1 | h-bond 2 | h-bond 1 | h-bond 2 | h-bond 1 | h-bond 2 | h-bond 1 | h-bond 2 |
| Closed | 0.98 | 0.57 | 0.96 | 0.11 | 0.85 | 0.51 | 0.18 | 0.03 |
| Open | 0.22 | 0.00 | 0.00 | 0.00 | 0.00 | 0.00 | 0.00 | 0.00 |

| | BAZ2B | | TRIM28 | | SMARCA2 | |
|---|---|---|---|---|---|---|
| | h-bond 1 | h-bond 2 | h-bond 1 | h-bond 2 | h-bond 1 | h-bond 2 |
| Closed | 0.93 | 0.28 | 0.94 | 0.54 | 0.82 | 0.59 |
| Open | 0.00 | 0.00 | 0.06 | 0.00 | 0.00 | 0.00 |

**Supplementary Table 4.** Percentage of frames that fulfill the geometrical criterion of a h-bond ($distance_{NH\cdots O}$ <= 0.25 nm and $angle_{NHO}$ >= 120º) for the two key h-bonds and the two metastable states of each BD. ZMYND11 is not included in the analysis because a proline (Pro199) is in place of the residue that acts as donor of h-bond 2.

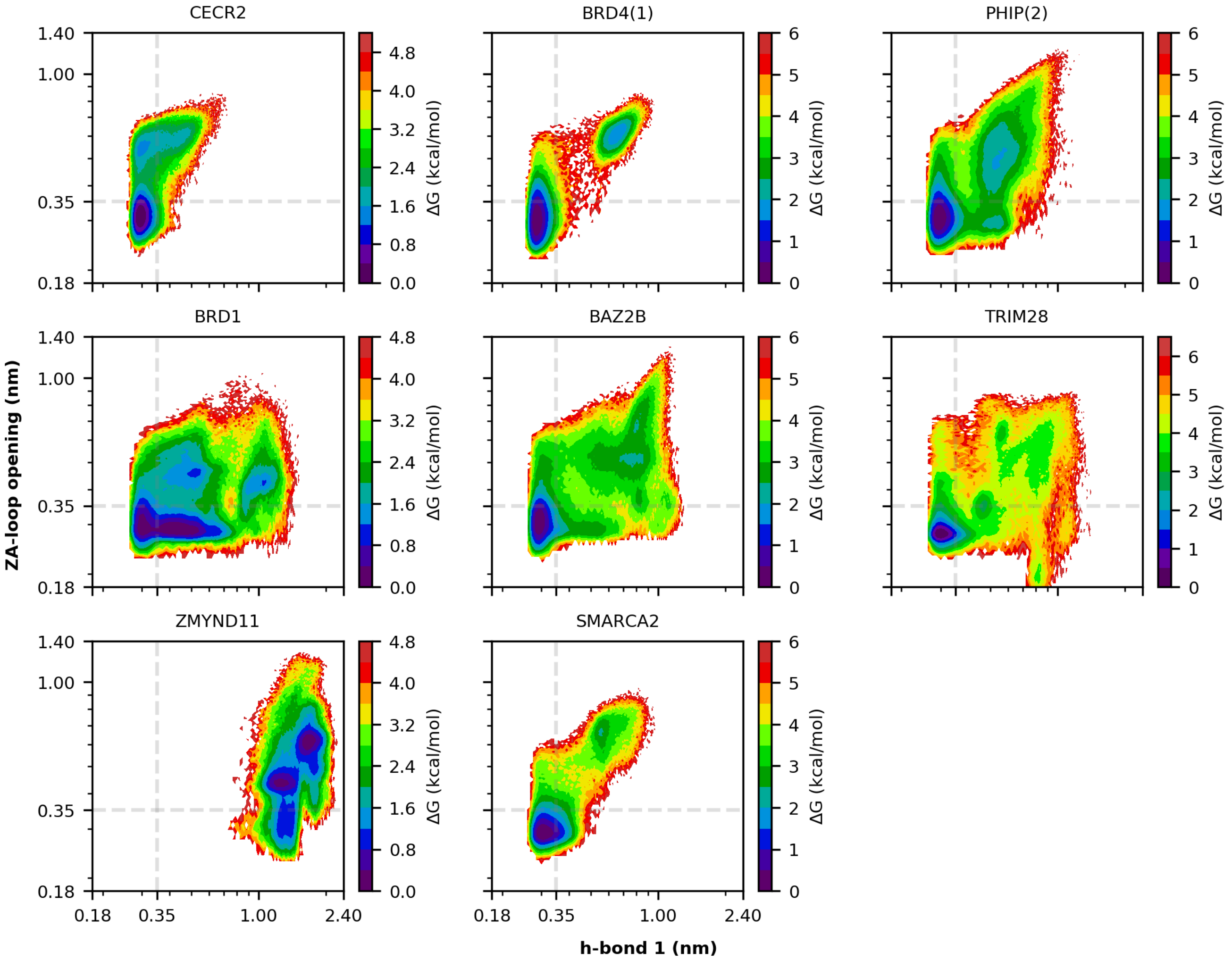


**Supplementary Figure 10. Raw free energy landscape of the key h-bond 1 and the ZA-loop opening for each BD.** Empirical free energy map computed via projection of all the simulation data onto h-bond 1 and the ZA-loop backbone contacts with the conserved aspartate. Axes are given in a logarithmic scale to facilitate comparison between BDs. Dashed lines indicate a distance of 0.35 nm as an upper bound for h-bond formation, which also represents one of the minimum possible contacts between the ZA-loop and the conserved aspartate.

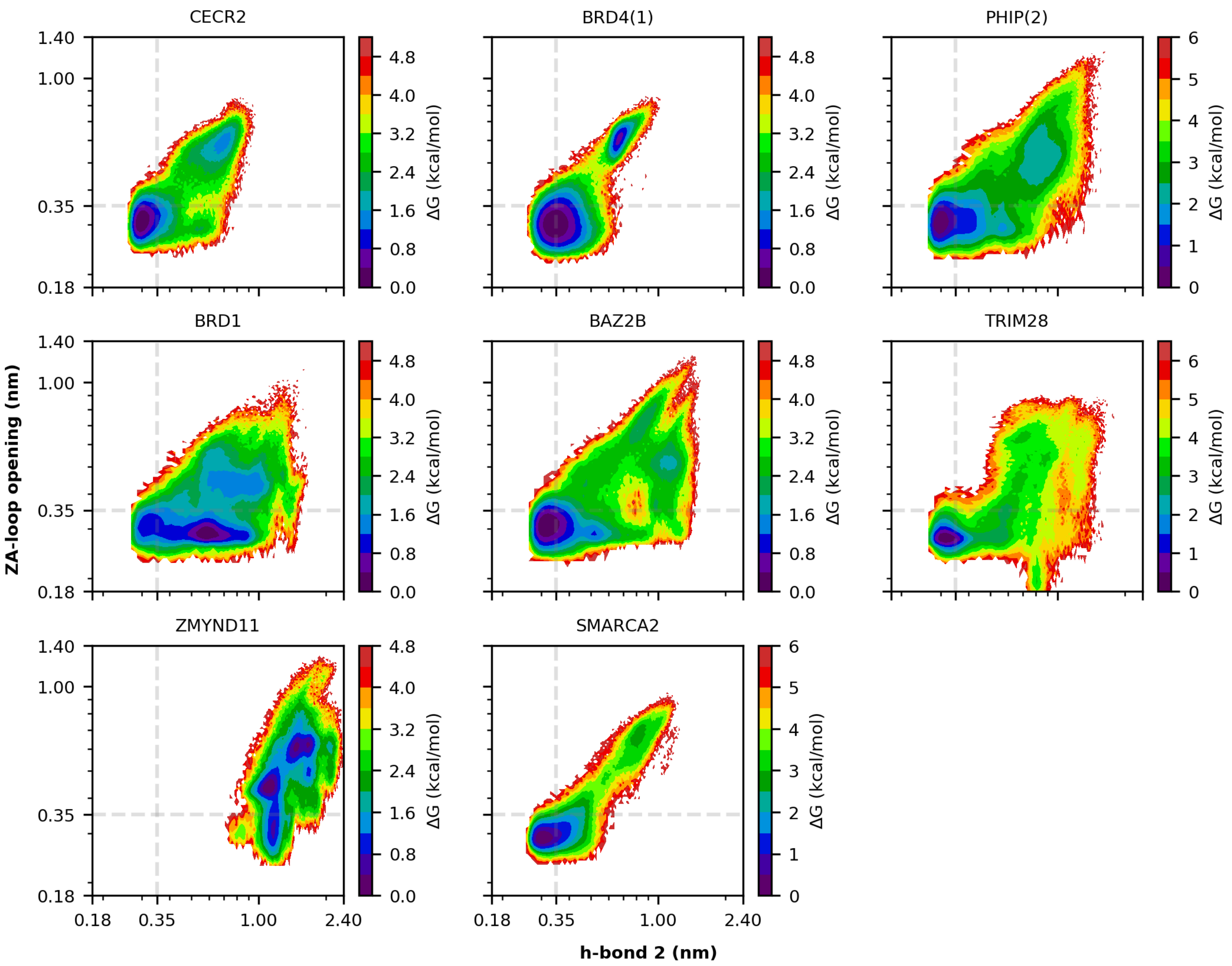


**Supplementary Figure 11. Raw free energy landscape of the key h-bond 2 and the ZA-loop opening for each BD.** Empirical free energy map computed via projection of all the simulation data onto h-bond 2 and the ZA-loop backbone contacts with the conserved aspartate. Axes are given in a logarithmic scale to facilitate comparison between BDs. Dashed lines indicate a distance of 0.35 nm as an upper bound for h-bond formation, which also represents one of the minimum possible contacts between the ZA-loop and the conserved aspartate.

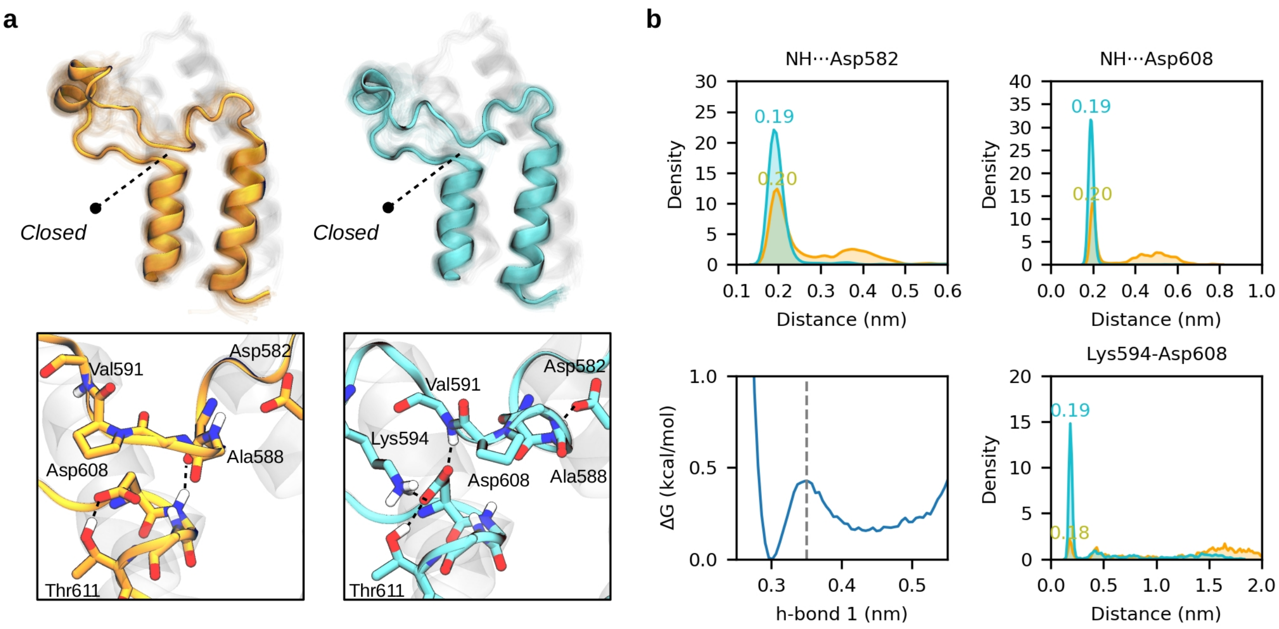


**Supplementary Figure 12. Alternative closed state in BRD1 where the ZA-loop slightly displaces towards the $\alpha_Z$ helix.** (**a**) Structure of the "closed" crystal-like state (orange) and the alternative closed state in which the ZA-loop is slightly displaced (cyan). A close view of the key region is represented below, highlighting the buried Asp582 and the conserved Asp608, both of which establish h-bonds with different backbone amides of the ZA-loop (mostly the ones of Ala588 and Val591). (**b**) Distribution of the minimum h-bond distance between selected ZA-loop amides and Asp582/Asp608, empirical free energy profile along h-bond 1, and minimum h-bond distance between Lys594 with Asp608 for the two metastable states. Note that the distributions in orange are bimodal and turn nearly unimodal (cyan) due to the conformational change.

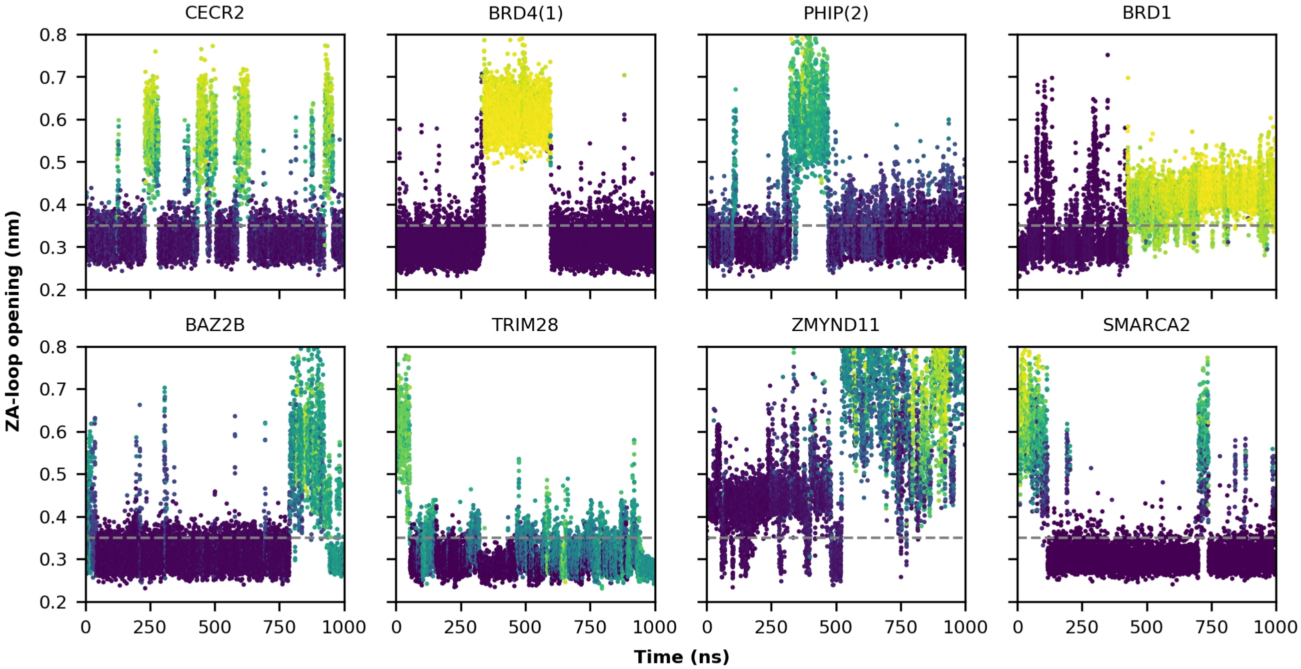


**Supplementary Figure 13. Selected trajectories showing sporadic openings of the ZA-loop for each BD.** Evolution of the ZA-loop opening contact distance along representative trajectories. Dots are colored according to their membership in the metastable open state characterized by each MSM, with green-yellow regions indicating a high probability and dark regions a low probability. Dashed lines indicate a distance of 0.35 nm as a reference of h-bond contacts. Note the sporadic, narrow and dark peaks that are present in all plots, indicating fast openings that lead to open states that are not metastable. Note as well that in ZMYND11 the closed state show contacts mostly above 0.35 nm, and is therefore only “semiclosed”.

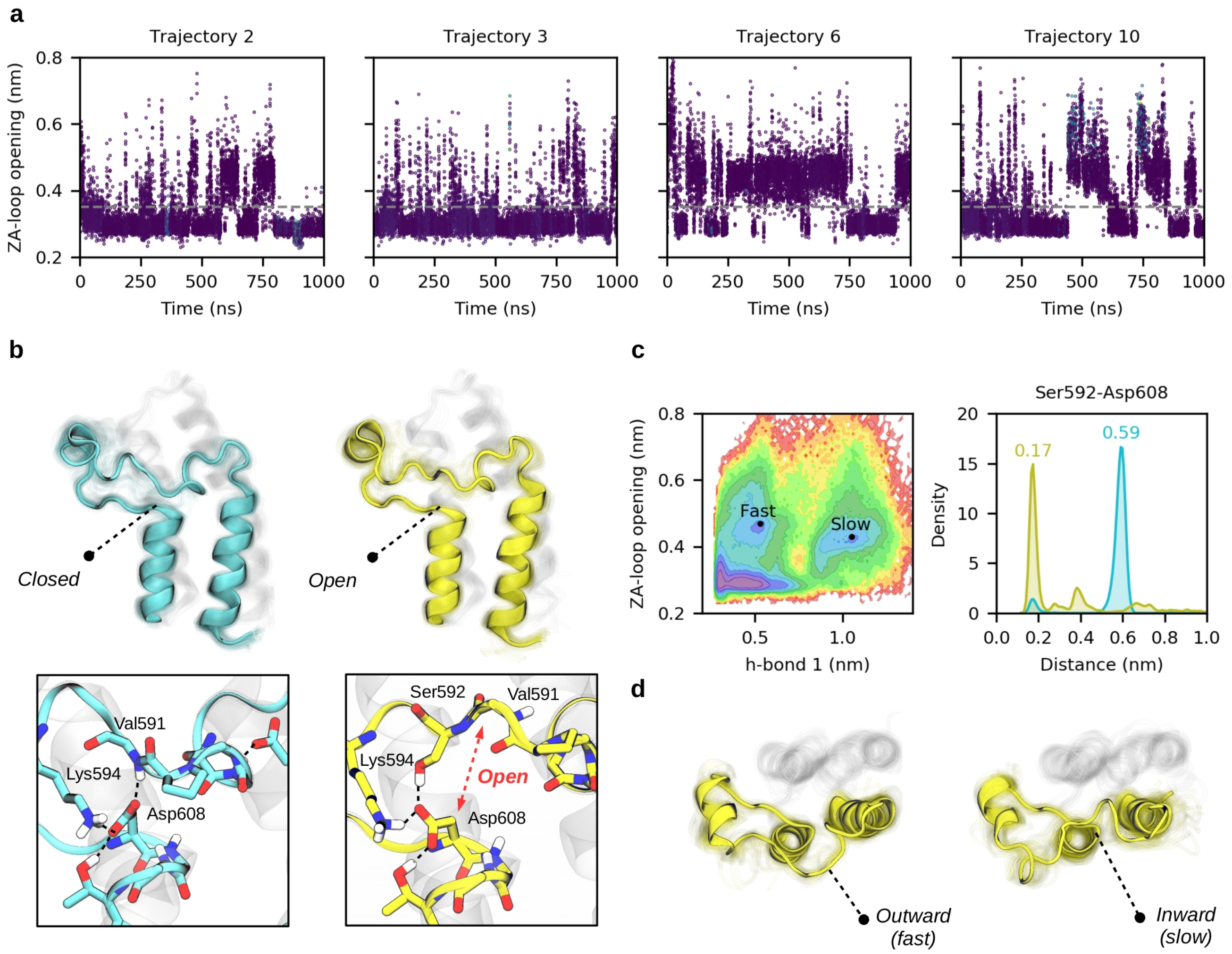


**Supplementary Figure 14. Selected trajectories showing sporadic openings of the ZA-loop in BRD1.** (**a**) Four different trajectories showing fast opening events, described by the ZA-loop opening contact distance. Dots have been colored according to their membership in the metastable open state characterized by the MSM, with black representing a low probability. Dashed lines indicate a distance of 0.35 nm as a reference of h-bond contacts. Note that for BRD1 the sporadic and narrow peaks are very frequent, and eventually show a certain degree of metastability. (**b**) The fast opening is mainly related to a switching interaction from the Val591-Asp608 backbone h-bond to the Ser592-Asp608 h-bond. (**c**) Empirical free energy landscape composed by h-bond 1 and the ZA-loop backbone contacts with the conserved aspartate. Distribution of the minimum distance between the side chains of Ser592 and Asp608 for the two states. (**d**) Top view of the fast/slow open state structures highlighting that the slow process is related to an outward-to-inward displacement of the ZA-loop, which is captured by the h-bond 1 distance.

#### 4. Analysis of pocket volumes

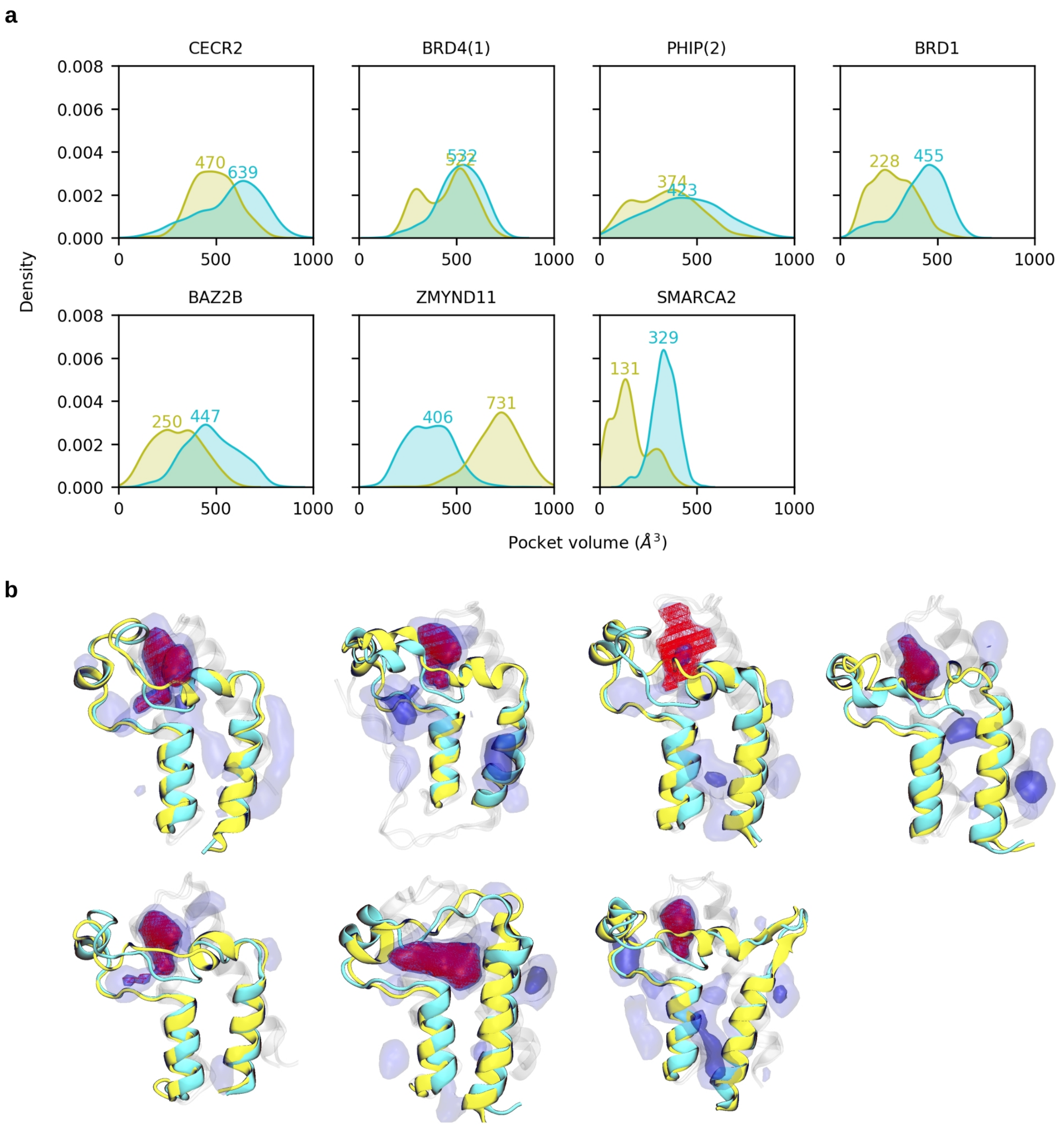


**Supplementary Figure 15. Shrinking of the acetyl-lysine pocket upon the opening process.** (**a**) Volume of the acetyl-lysine pocket of each BD evaluated with 1.000 structures of the closed state (cyan) and the open state (yellow). The mode of each kernel density estimate is shown as a colored number. (**b**) Visual representation of the acetyl-lysine pockets –shown in red– and the two metastable states for each BD. Note that for ZMYND11 we selected the principal cryptic pocket since the closed state is never fully formed and thus the acetyl-lysine pocket is not defined. TRIM28 is not included in this analysis given the lack of noticeable pockets in the closed state.

| | CECR2 | | BRD4(1) | | PHIP(2) | | BRD1 | |
|---|---|---|---|---|---|---|---|---|
| | Open | Closed | Open | Closed | Open | Closed | Open | Closed |
| Samples | 999 | 997 | 999 | 997 | 967 | 896 | 930 | 974 |
| Mean | 493 | 574 | 451 | 517 | 339 | 437 | 272 | 413 |
| SD | 114 | 166 | 127 | 115 | 160 | 193 | 110 | 128 |
| Min | 75 | 29 | 163 | 103 | 27 | 25 | 72 | 46 |
| 25% | 409 | 458 | 328 | 447 | 206 | 300 | 186 | 344 |
| Median | 488 | 604 | 478 | 527 | 343 | 435 | 263 | 438 |
| 75% | 572 | 697 | 548 | 599 | 456 | 577 | 355 | 509 |
| Max | 815 | 973 | 739 | 779 | 793 | 971 | 601 | 675 |

| | BAZ2B | | ZMYND11 | | SMARCA2 | |
|---|---|---|---|---|---|---|
| | Open | Closed | Open | Closed | Open | Closed |
| Samples | 912 | 995 | 997 | 973 | 524 | 991 |
| Mean | 303 | 478 | 711 | 362 | 157 | 334 |
| SD | 124 | 135 | 121 | 118 | 96 | 65 |
| Min | 32 | 101 | 173 | 57 | 25 | 113 |
| 25% | 205 | 384 | 637 | 271 | 92 | 296 |
| Median | 296 | 470 | 721 | 363 | 140 | 335 |
| 75% | 395 | 575 | 791 | 446 | 210 | 378 |
| Max | 623 | 846 | 1034 | 839 | 428 | 540 |

**Supplementary Table 5.** Volume distribution of the acetyl-lysine pocket ($Å^3$) of each BD evaluated with 1000 representative structures of the open and closed states. The number of samples correspond to volumes above 0 $Å^3$. Volumes for ZMYND11 refer to the principal cryptic pocket since the acetyl-lysine pocket is not defined. Note the high standard deviation (SD) of the pocket volume of PHIP(2) in the closed state. TRIM28 is not included in this analysis given the lack of noticeable pockets in the closed state.

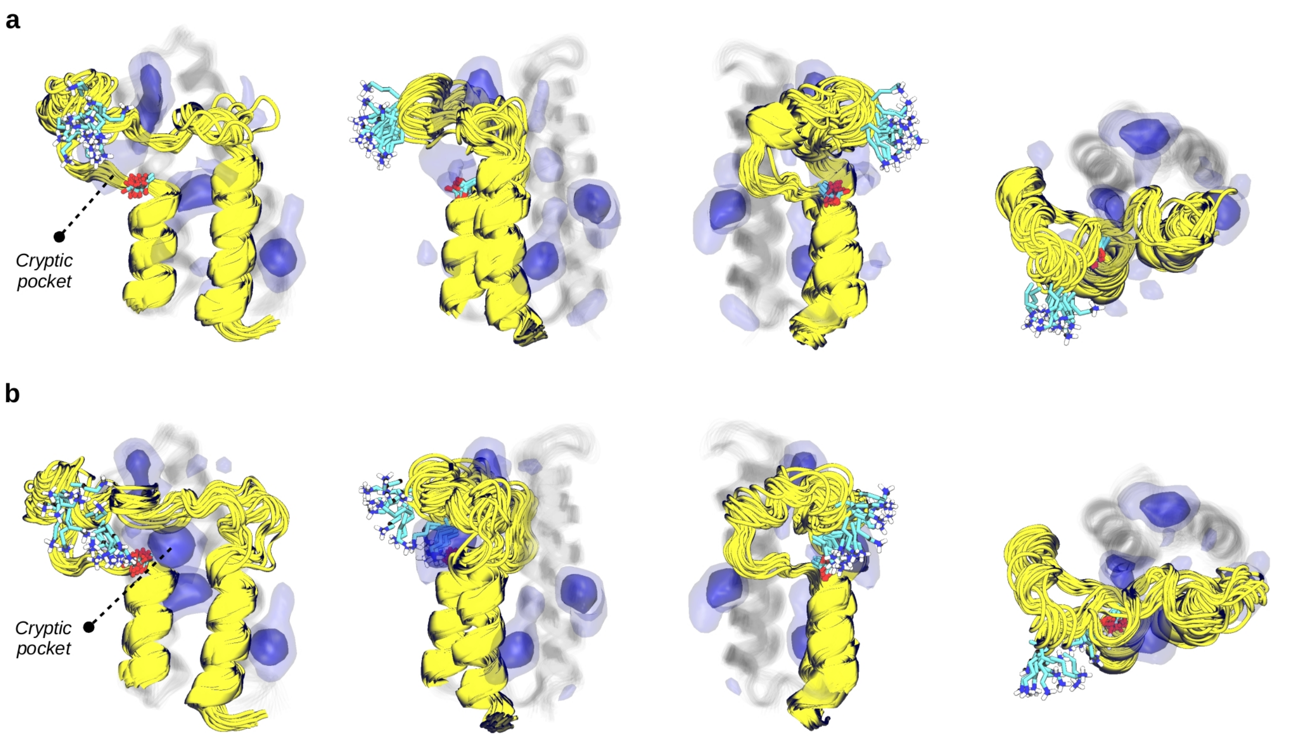


**Supplementary Figure 16. Displacement of the cryptic pocket in an alternative open state of BRD1.** Different views of the cryptic pocket in the open state characterized by the MSM (**a**) and in an alternative open state that is not included in the model (**b**). Note that in the alternative state the pocket is displaced towards the $\alpha_Z$ and $\alpha_A$ helices and that a lysine residue (Lys594, shown in cyan) interacts with the conserved aspartate (Asp608). Pocket frequency maps are represented by blue isosurfaces given at 0.25 (light) and 0.50 (intense) isovalues.

#### 5. Analysis of experimental structures

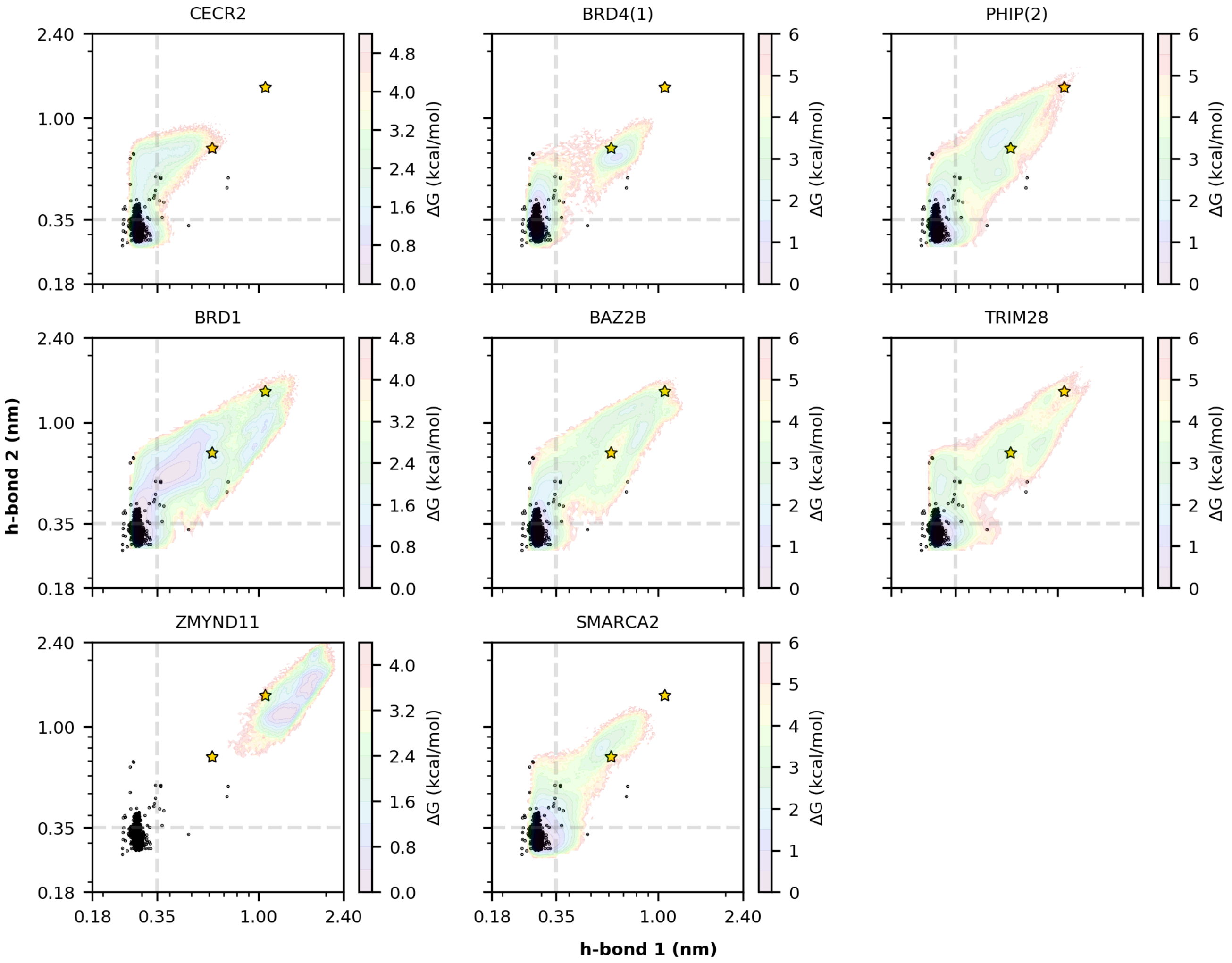


**Supplementary Figure 17. Distribution of the key h-bonds in experimental structures projected onto the raw h-bond free energy landscape of each BD.** All experimental structures (black dots) are obtained from the Pfam database (PF00439). The two stars indicate the crystal structures of PB1(6) and ZMYND11 (3IU6 and 4N4G) that are found in an open state. Empirical free energy maps are computed via projection of all the simulation data onto h-bond 1 and h-bond 2. Axes are given in a logarithmic scale to facilitate comparison between BDs. Dashed lines indicate a distance of 0.35 nm as an upper bound for h-bond formation. Note that there is a remarkable agreement between the distribution of experimental structures and the regions explored by our MD simulations. Note as well that ZMYND11 is an exception since it is stable in the open state.

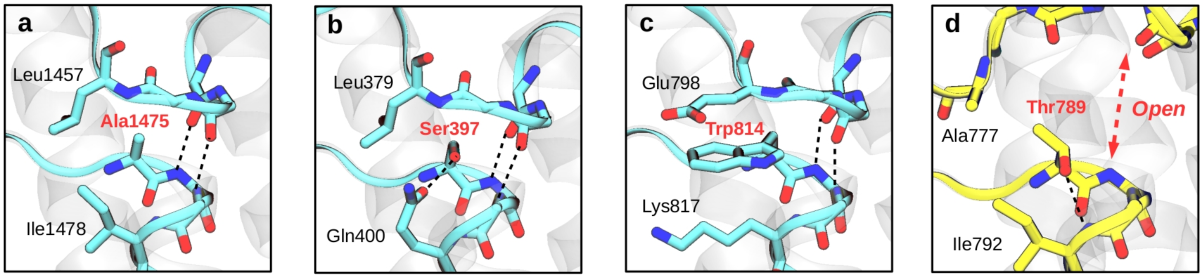


**Supplementary Figure 18. Closeup of four experimental structures of BDs that do not display the conserved aspartate.** (**a**) Structure of BAZ1A (PDB 5UIY) with Ala1475 in place of the conserved aspartate. Note the presence of hydrophobic residues Leu1457 and Ile1478 surrounding the alanine; (**b**) Structure of PB1(3) (PDB 3K2J) with Ser397 in place of the conserved aspartate. Note that Gln400 establishes an h-bond with the serine; (**c**) Structure of SP100 (PDB 4PTB) with Trp814 in place of the conserved aspartate. Note the presence of charged residues Glu798 and Lys817 that could establish π-interactions with the tryptophan. (**d**) Structure of PB1(6) (PDB 3IU6) with Thr789 in place of the conserved aspartate. Hydrophobic residues Ala777 and Ile792 are surrounding the threonine, which is also hydrogen bonded with the amide group of Ile792. Note that in this case the two backbone h-bonds are not formed and the ZA channel is disrupted, likely because of the bulky methyl group that threonine has in its beta carbon.

#### 6. Nuclear magnetic resonance predictions

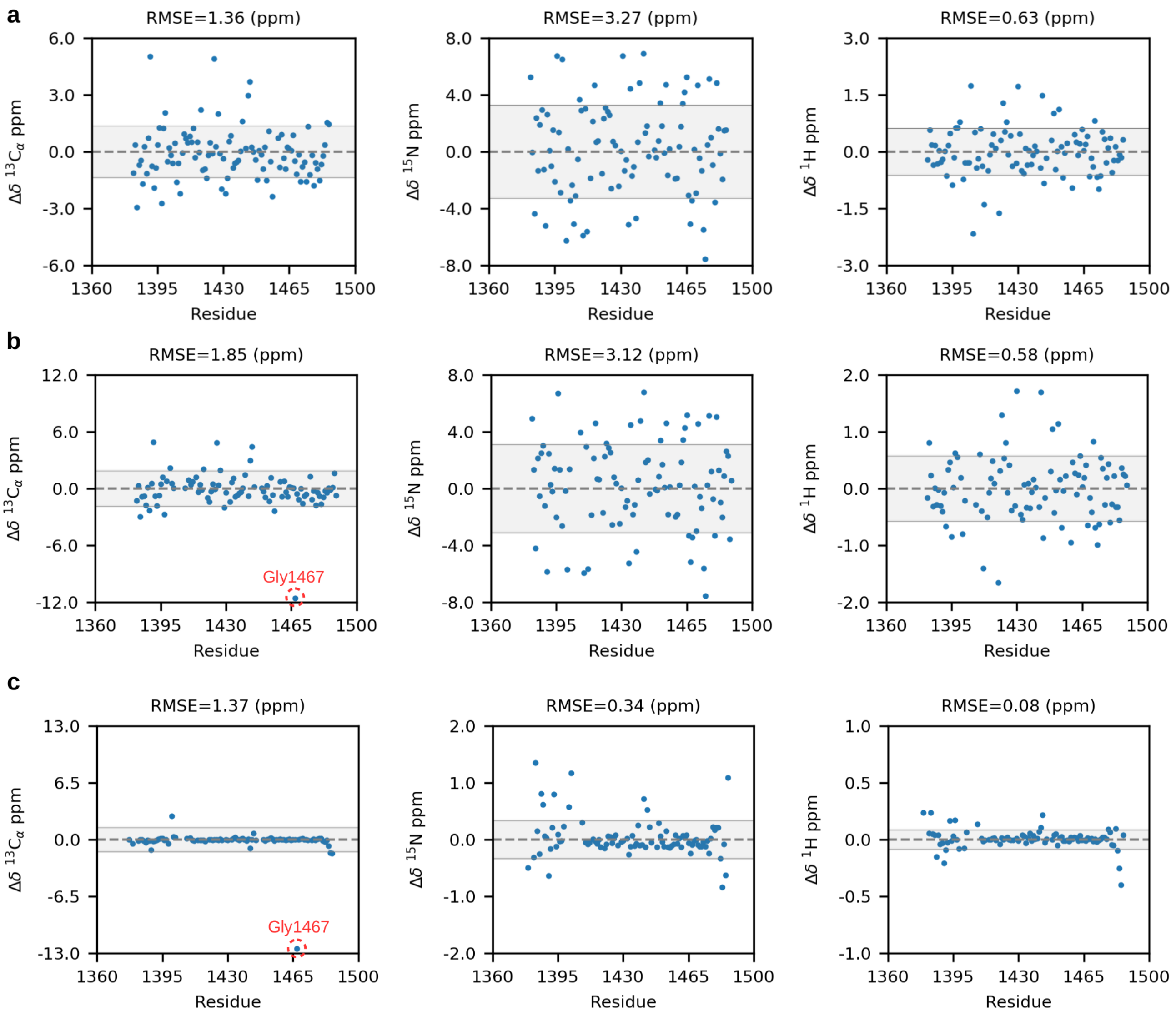


**Supplementary Figure 19. Comparison of NMR SMARCA2 expectation values and experimental data.** Difference between $^{13}$C, $^{15}$N and $^{1}$H CAMSHIFT MSM expectation values and the experimental values of BMRB 11329 (**a**) and BMRB 27106 (**b**). The last panel (**c**) compares the two experimental sources to highlight the large deviation of Gly1467 in the $^{13}$C plot, which may arise from a misassignment or because in BMRB 27106 the hairpin has 18 extra amino acids that could interact with the glycine. The shadow areas indicate root-mean square errors (RMSE) and the residue numbering is that of PDB 5DKC.

| | $^{13}C_{\alpha}$ | | $^{15}N$ | | $^{1}H$ | |
|---|---|---|---|---|---|---|
| **a** | Residue | Δδ (ppm) | Residue | Δδ (ppm) | Residue | Δδ (ppm) |
| 1 | ILE1391 | 5.05 | ILE1475 | -7.55 | SER1406 | -2.17 |
| 2 | ILE1425 | 4.91 | LYS1442 | 6.93 | LEU1405 | 1.75 |
| 3 | ARG1444 | 3.71 | PHE1431 | 6.77 | ASP1430 | 1.73 |
| 4 | TYR1443 | 2.96 | ASN1396 | 6.74 | GLU1420 | -1.62 |
| 5 | THR1384 | -2.93 | ASP1399 | 6.52 | TYR1443 | 1.48 |
| 6 | TYR1397 | -2.73 | SER1401 | -6.28 | LEU1412 | -1.40 |
| 7 | LEU1456 | -2.35 | ILE1410 | -5.90 | TYR1422 | 1.29 |
| 8 | GLU1407 | -2.21 | LEU1412 | -5.62 | ASP1452 | 1.13 |
| 9 | LEU1418 | 2.21 | SER1474 | -5.49 | LEU1449 | 1.01 |
| 10 | PHE1431 | -2.20 | LYS1382 | 5.25 | ASP1473 | -0.98 |
| **b** | Residue | Δδ (ppm) | Residue | Δδ (ppm) | Residue | Δδ (ppm) |
| 1 | GLY1467 | -11.61 | ILE1475 | -7.57 | ASP1430 | 1.72 |
| 2 | ILE1391 | 4.89 | LYS1442 | 6.80 | TYR1443 | 1.70 |
| 3 | ILE1425 | 4.86 | ASN1396 | 6.71 | GLU1420 | -1.66 |
| 4 | ARG1444 | 4.44 | ILE1410 | -5.95 | LEU1412 | -1.41 |
| 5 | TYR1443 | 2.97 | ILE1390 | -5.85 | TYR1422 | 1.29 |
| 6 | THR1384 | -2.95 | SER1401 | -5.70 | ASP1452 | 1.15 |
| 7 | TYR1397 | -2.75 | LEU1412 | -5.68 | LEU1449 | 1.05 |
| 8 | LEU1456 | -2.36 | SER1474 | -5.62 | ASP1473 | -0.99 |
| 9 | ALA1389 | -2.33 | ILE1434 | -5.27 | ASN1459 | -0.95 |
| 10 | SER1400 | 2.19 | GLY1467 | -5.19 | ARG1444 | -0.87 |
| **c** | Residue | Δδ (ppm) | Residue | Δδ (ppm) | Residue | Δδ (ppm) |
| 1 | GLY1467 | -12.49 | LEU1383 | 1.36 | ARG1485 | -0.40 |
| 2 | SER1400 | 2.71 | GLY1402 | 1.18 | ALA1484 | -0.25 |
| 3 | GLN1486 | -1.58 | GLN1486 | 1.10 | ASN1379 | 0.24 |
| 4 | ARG1485 | -1.50 | SER1483 | -0.84 | LEU1383 | 0.24 |
| 5 | ALA1389 | -1.18 | GLN1386 | 0.81 | TYR1443 | 0.22 |
| 6 | LYS1442 | -0.96 | THR1393 | 0.79 | ILE1390 | -0.21 |
| 7 | ALA1484 | -0.75 | HIS1441 | 0.72 | ASN1396 | 0.17 |
| 8 | ARG1444 | 0.73 | ILE1390 | -0.64 | THR1393 | 0.17 |
| 9 | ASN1379 | -0.48 | ARG1485 | -0.62 | GLN1386 | -0.15 |
| 10 | LYS1398 | -0.47 | MET1387 | 0.62 | VAL1408 | 0.14 |

**Supplementary Table 6.** Largest deviations between $^{13}C$, $^{15}N$ and $^{1}H$ CAMSHIFT MSM expectation values and the experimental values of BMRB 11329 (**a**), BMRB 27106 (**b**) and the comparison of the two experimental sources (**c**). The residue numbering is that of PDB 5DKC.
